# TGA family of transcription factors are nitric oxide sensors in the root stem cell niche

**DOI:** 10.1101/2025.10.14.682262

**Authors:** Sara Gómez-Jiménez, María Guadalupe Fernández-Espinosa, Capilla Mata-Pérez, Álvaro Sanchez-Corrionero, Jina Song, María Angels de Luis Balaguer, Rosangela Sozzani, Nora Gutsche, Sabine Zachgo, Jelle Van Leene, Geert De Jaeger, Steven H. Spoel, Oscar Lorenzo

## Abstract

Plant root development is tightly regulated and depends on the coordination among different cell types located in the root stem cell niche (SCN). A complex network of regulators includes the gasotransmitter nitric oxide (NO) known to inhibit primary root growth, modify root architecture and alter meristem organisation. However, little is known about the molecular targets of this signalling molecule during root SCN homeostasis. Here, genetic, molecular and physiological evidence support that TGA transcription factors (TFs) of the bZIP family, and specifically PERIANTHIA (PAN), contribute to the regulation of the root SCN through NO sensing. This gasotransmitter induces reversible S-nitrosylation of PAN, thereby inhibiting its binding to DNA and associated transcriptional activation; whilst the protein-SNO reductase activity of TRXh5 can denitrosylate PAN. This switch on and off mechanism for PAN activity modulates gene expression in the SCN and the functioning of the quiescent centre, thereby altering columella stem cell maintenance through a NO-dependent mechanism. Furthermore, PAN interacts selectively with several TGA members which are susceptible to differential S-nitrosylation. Given the functional redundancy within the TGA family, analysis of quadruple and pentuple mutants revealed close coordination between PAN and other members of this family of TFs, which contributes to the NO sensing during the development of the root apical meristem. Collectively, these findings reveal a molecular framework in which the TGA family, primarily coordinated by PAN activity, acts as a master NO sensor in the root SCN.

## Introduction

Root development is a tightly regulated process that depends on coordination among the different cell types located in the stem cell niche (SCN). The latter includes a group of mitotically less-active cells, known as quiescent centre (QC), which is surrounded by the initial stem cells that acquire different cell fates through their division, to give rise to different root tissues such as columella cells (Cruz-Ramírez et al. 2012). The maintenance of the SCN involves a complex network of regulatory factors among which is the gasotransmitter nitric oxide (NO) (Fernández-Marcos et al. 2011; Sanz et al. 2014). NO is a ubiquitous signalling molecule involved in the regulation of many different processes throughout plant development and stress compensation (reviewed in Sanz et al., 2015; Sánchez-Vicente et al., 2019; Sanchez-Corrionero et al., 2022) NO has been shown to modulate primary root (PR) growth, the organization of the root apical meristem (RAM) and stem cell decisions, affecting the size and number of meristem cells (Fernández-Marcos et al. 2011, 2012; Sanz et al. 2014). NO perception and signal transduction occur mainly through posttranslational modifications (PTMs) of target proteins, altering their activity, localization and/or stability (Hess et al. 2001; Astier and Lindermayr 2012; Kovacs and Lindermayr 2013; Mata-Pérez et al. 2023). *S*-nitrosylation of cysteine residues stands out as the most relevant NO-PTM (Hess et al. 2001; Astier and Lindermayr 2012; Kovacs and Lindermayr 2013; Mata-Pérez et al. 2023). This dynamic and reversible modification consists of the covalent binding of NO to a thiol group of a cysteine residue of the target protein within a suitable microenvironment, generating an *S*-nitrosothiol group (-SNO).

Certain members of the basic (region) leucine zipper (bZIP) transcription factor (TF) family of *Arabidopsis thaliana* (Arabidopsis) are an example of proteins subjected to these alterations (Albertos et al. 2015; Lorenzo 2019). Among them, the TGA family (according to its conserved DNA-binding motif TGACG) comprises prototypical members with functional and regulatory relevance. For instance, the redox regulation of TGA1, a member of clade I, involves the *S*-nitrosylation at several cysteine residues (Lindermayr et al. 2010). Similarly, the TGA family member PERIANTHIA (PAN) has been reported to undergo redox-sensitive DNA-binding with decreased binding observed under oxidising conditions. This sensitivity is mediated by N-terminal cysteines Cys68 and Cys87 that can form a disulfide bond, while Cys340 in the C-terminal part can be *S*-glutathionylated, which is a requirement for PAN activity (Li et al. 2009; Gutsche and Zachgo 2016). In addition to the involvement of PAN in floral development (Chuang et al. 1999) and SAM organisation (Maier et al. 2009, 2011), it has also been identified as an important molecular regulator of root SCN and, more specifically, of quiescent centre (QC) function and columella stem cell (CSC) maintenance (Hu et al. 2022).

Some TGA members have been linked to functions in the root. For example, TGA1 and TGA4 are involved in root architecture (Alvarez et al. 2014; Canales et al. 2017), and there is functional redundancy between TGAs during root growth (Hu et al. 2022) with overrepresentation of TGA *cis*-regulatory elements in genes active in root tip cells (Van Der Zaal et al. 1991). However, the specific role of the TGA group in controlling root SCN remains unknown. Here we unveil the regulatory mechanisms of root growth and stem cell patterning by NO and provide a molecular framework for the involvement of PAN and other TGA family members in root development, stem cell maintenance and NO sensing in this vital plant tissue.

## Material and methods

### Plant material

*Arabidopsis thaliana* Columbia-0 ecotype (Col-0) was the genetic background used in this work. The *pan051790* seeds were obtained from ABRC (SALK_057190), *pPAN:GFP* (ABRC; Lee et al. 2006) and *XVE:PAN* from the TRANSPLANTA collection (Coego et al. 2014). The single *tga7,* triple *tga2-1 tga5-1 tga6-1* and quadruple *tga2-1 tga3-1 tga5-1 tga6-1* Arabidopsis mutants were kindly provided by Dr. Pablo Tornero (IBMCP-CSIC, Valencia, Spain).

### Growth conditions

For *in vitro* culture, Arabidopsis seeds were surface-sterilised in 75% (v/v) household bleach and 0.01% (v/v) Triton X-100 for 5 min and washed three times in sterile water before sowing. Seeds were stratified for 3 days at 4°C and then sown on Murashige and Skoog (MS) solid medium with MES (0.23% (w/v)), 0.75% (w/v) sucrose and 1.5% (w/v) agar, and the pH was adjusted to 5.8 with KOH before autoclaving. Seeds were sown on the media in the presence or absence of different treatments (1 mM cPTIO, 0.5 mM GSNO, 0.5 mM SNAP) and vertical plates were sealed and incubated in a controlled environment growth chamber (22°C, 60% relative humidity, and a long-day photoperiod).

### Generation of expression vectors

Full-length cDNAs were amplified with first strand cDNA synthesis system Superscript (ThermoFisher Scientific) from RNA obtained from Arabidopsis seedlings extracted with TRIzol reagent (ThermoFisher Scientific). The coding sequences (CDS) of *PAN*, *TGA2*, *TGA3* and *TGA5* were amplified with primers indicated in Supplementary Table 3 and *Taq* polymerase (Roche) by PCR (Sambrook et al. 1989). PCR products were purified with GeneClean kit (MP Biomedicals) and then cloned into gateway entry clone pENTR/D-TOPO (ThermoFisher Scientific). They were then transferred to pYFPC43 and pYFPN43 vectors (Curtis and Grossniklaus 2003; Belda-Palazón et al. 2012) for transient expression and to pDEST17 (ThermoFisher Scientific) vector for protein expression and purification. Sequence accuracy was confirmed by sequencing before delivered into *Escherichia coli* TOP10 strain (Invitrogen) and *Agrobacterium tumefaciens C58C1* strain with pGV2260 virulence plasmid (Deblaere et al. 1985).

### Knock-out mutant isolation and generation of multiple mutants

*pan* SALK mutant line *051790*, the triple *tga2-1tga5-1tga6-1* and quadruple *tga2-1tga3-1tga5-1tga6-1* mutants were genotyped from plant genomic DNA. Genomic DNA was obtained by CTAB DNA extraction method (J. 1990). DNA was amplified with primers in Supplementary Table 3 and *Taq* Polymerase (Roche) by PCR (Sambrook et al. 1989).

*pantga2tga3tga5tga6* mutant was generated by crosses. Homozygous lines were identified by PCR genotyping of segregating F2 populations with oligonucleotides described in Supplementary Table 3.

### Site-directed mutagenesis

Site-directed mutagenesis of *PAN* was performed using the QuickChange II Site Directed Mutagenesis Kits (Stratagene Corporate). Plasmid pDEST17 with CDS of PAN was used as template and primers were designed using the tools from Stratagene and synthesised by Metabiom. The primers used are listed in Supplementary Table 3. Mutations were confirmed by sequencing (Sequencing Facility, NUCLEUS, University of Salamanca).

### Subcellular localization in *Nicotiana benthamiana*

Coding sequences of WT *PAN* and cysteine 340-to-serine (*C340S*) mutant version were cloned in pGWB6 (*35S:GFP-PAN* and *35S:GFP-PANC340S*) vector (Nakagawa et al. 2009). The constructs were transiently expressed into *Nicotiana benthamiana* epidermal cells by agroinfiltration of transformed *Agrobacterium tumefaciens* C58C1 with pGV2260 (Deblaere et al. 1985). Additionally, p19 was added to avoid silencing. Leaves were observed by confocal microscopy after 72 hours. GFP was excited with the 488 nm argon laser and detection filters between 515-530 nm in a Leica TCS SP2 confocal microscope.

### Bimolecular fluorescence complementation assay

The coding regions of *PAN*, *TGA2*, *TGA3* and *TGA5* genes were fused to the *C*-terminal end of the YFP in the pYFC43 and pYFN43 (Curtis and Grossniklaus 2003; Belda-Palazón et al. 2012). YFP-fusions and control constructs were then transiently expressed into *Nicotiana benthamiana* epidermal cells by agroinfiltration of transformed *Agrobacterium tumefaciens* C58C1 with pGV2260 (Deblaere et al. 1985) and p19 to avoid silencing. Leaves were observed by confocal microscopy after 72 hrs of incubation. For live imaging of YFP, fluorescence was examined with a Leica TCS SP2 confocal microscope.

### Measurement of primary root length and cell size

Arabidopsis plants were grown for 7 days in MS media supplemented with 5 µM *β*-estradiol, 5 µM *β*-estradiol combined with 0.5 mM SNAP or 5 µM *β*-estradiol combined with 1 mM cPTIO, and seedlings were scanned at 600 dots per inch. Root length was measured using ImageJ software (Schindelin et al. 2012). Statistical comparisons were performed using the test appropriate for each experimental design, as indicated in the corresponding figure legends, with *P*-values adjusted using the Benjamini–Hochberg procedure for multiple comparisons.

To measure root cell size, seedlings were stained with the modified mPS-PI protocol (Truernit et al. 2008). Confocal stacks of roots were obtained with a Leica SP 2 confocal microscope and cell length was measured using ImageJ software. Up to 30 consecutives cortex cells were measured per root and grouped according to their position from the QC as cells 1–10, 11–20 and 21– 30. Cell-size data were analysed using a three-way aligned rank transform (ART) ANOVA with genotype, treatment and cell group as fixed factors and root identity as a random effect. Following a significant three-way interaction, pairwise comparisons among treatments were performed within each genotype and cell group using estimated marginal means, with *P*-values adjusted using the Benjamini–Hochberg false discovery rate procedure.

### Fluorescence microscopy analysis

Arabidopsis plants were grown for 5 days on a mesh vertical plate and transferred to MS medium containing 1 mM cPTIO or 0.5 mM SNAP. Staining of root cells with propidium iodide for the GFP analysis of *pPAN:GFP* plants was performed using confocal laser scanning microscopy Zeiss LSM 710. The excitation source was an argon ion laser at 488 nm, and detection filters were between 515 and 530 nm. To quantify fluorescence from confocal images, the laser power and detector gain were adjusted so that the sample with the higher fluorescence intensity did not contain saturated pixels. Fluorescence was quantified using Fiji, an open-source image processing package based on software ImageJ. The region of interest (ROI) was hand-drawn in each image, and the area, integrated density and mean grey value were measured. The background was subtracted from the ROI by defining a rectangular ROI with a fixed area in an area that did not contain fluorescence and adjacent to the initial ROI. This step was repeated three times, and the average mean grey value of the background was used to calculate the corrected total cell fluorescence (CTCF), according to the formula: CTCF = Integrated density – (Area * Mean grey value of background ROI).

### Gravitropic response

Seedlings from Col-0, *pan051790*, triple *tga2tga3tga5tga6*, quadruple *tga2tga3tga5tga6* and pentuple *pantga2tga3tga5tga6* genotypes were grown vertically as described above. Four days post-imbibition (4 dpi), germinated seedlings were transferred to a fresh plate and one day after (5 dpi) they were turned 90 degrees as described elsewhere (Sato et al. 2015). Plates were scanned 6h after gravistimulation and root angles were measured using ImageJ software. A minimum of three replicates were used in this study. Data normality was assessed using the Shapiro–Wilk test and homogeneity of variances using Levene’s test. As the data did not meet the assumptions of normality and homoscedasticity, differences among genotypes were analysed using the Kruskal–Wallis test. As significant differences were detected, pairwise comparisons were performed using Dunn’s post hoc test with Benjamini– Hochberg adjustment for multiple comparisons. Differences were considered statistically significant at *P* < 0.05.

### RNA-Seq differential expression analysis

For RNA-Seq purposes, 5-day-old *XVE:PAN* Arabidopsis roots were dissected at approximately 0.1 cm from the root tip. Before the end of the 5 days, plants were treated during 6 and 24 hrs with 5 μM *β*-estradiol, 500 μM SNAP, 1 mM cPTIO and combinations of *β*-estradiol with SNAP or cPTIO. Three biological replicates of each condition were assessed. RNA was extracted using the RNeasy Micro Kit (Qiagen). cDNA synthesis, amplification and library preparation were performed using the NEBNext Poly(A) mRNA Magnetic Isolation Module and the NEBNext Ultra RNA Library Prep Kit for Illumina (NEB). Libraries were sequenced on an Illumina HiSeq 2500 with 100 bp single end reads in the Genomic Sciences Laboratory in North Carolina State University.

FastqQC version 0.11.9 was used to analyse reads quality (Andrews 2010). Reads were mapped against the TAIR 10 Reference Genome to quantify the expression in TPM values HISAT2 version 2.1.0 (Kim et al. 2019). Differential expression analysis was done with DESeq2 algorithm (Love et al. 2014) and DEGs were selected based on q-value ≤ 0.05 and fold-change ≥ |1.5| from the comparison of treated to untreated samples. Volcano and scatter plots were generated with ggplot2 (Wickham 2016) and, Venn Diagrams with ggVennDiagram R packages (Gao and Dusa 2024) and heatmaps were generated with pheatmap package (Kolde 2019).

For the heatmap analysis, up- and down-regulated genes (711 in total after *β*-estradiol and cPTIO treatment) were obtained from Venn diagrams while expression values (FPKM) came from the root map (Li et al. 2016) using Microsoft Access through a query. The query was the combination of the gene IDs, fold change and expression values (Li et al. 2016) through TAIR.

### Production of recombinant proteins and polyclonal antibodies

To produce PAN protein, wild type (WT) version of PAN recombinant protein was expressed in *Escherichia coli* (Biomedal). The bacterial culture was induced at different conditions and the soluble and insoluble fractions were analysed. Protein extraction was carried out by using a denaturing buffer containing 8 M urea, followed by on-column refolding (by reducing the urea concentration) and elution was performed using an imidazole gradient (10 mM to 1 M) added to the binding buffer without urea from the IMAC resin (High density nickel resin, ABT). Final protein was solubilized in 20 mM Tris-HCl pH 8.0, 3.8 mM DTT, and 0.2% SDS to avoid precipitation and analysed by western blot using an anti-His antibody (Genescript). Polyclonal PAN antibody production was carried out in rabbits as described in Albertos et al. (2015)(Albertos et al. 2015).

Wild-type and mutated PAN recombinant proteins were expressed in *Escherichia coli* BL21 (DE3) pLysS strain for protein amplification by induction with isopropyl *β*-D-1-thiogalactopyranoside (IPTG) and purified using BugBuster kit (Novagen) by Ni-NTA His Bind Resin (Novagen) according to the manufacturer’s protocol.

### Wheat germ protein production

*In vitro*-translated epitope-tagged proteins were produced in wheat germ extracts by using the Next Generation Cell Free Protein Expression Kit (Wheat Germ, Sigma-Aldrich). TGA1 (AT5G62510), TGA2 (AT5G06950), TGA5 (AT5G06960) and TGA7 (AT1G77920) were obtained from RIKEN Arabidopsis full-length cDNA clones (RAFL clones) developed by RIKEN Genomic Sciences Center (GSC) in Yokohama, Japan. TGA3 (AT1G22070) was cloned into pDONR221, TGA6 (AT3G12250) was obtained from SIGnAL (Salk pUni Clone U87058) and TGA8 (AT1G68640) was cloned into pCR8/GW/TOPO using the Gateway cloning system (ThermoFisher). Protein synthesis was performed using the *in vitro* Transcription/Translation Reagents according to manufacturer’s instructions (Sigma-Aldrich). For in vitro transcription, the coding DNA sequence of FLAG tag (DYKDDDDK) was attached to the cDNA templates of transcription factors by PrimeStar polymerase (Takara). The RNA samples were dissolved in 70 μl of RNase free water and mixed with 20 μl of a wheat germ extract and 20 μl of amino acid mixture at 16°C overnight. The synthesised protein was confirmed by western blotting.

### Western blotting

Total protein extraction for western blot purposes was carried out using 5- or 7-day-old Arabidopsis seedlings. Tissue was ground in liquid nitrogen using glass beads and SILAMAT mixer and homogenised in 1 volume of extraction buffer (100 mM Tris-HCl, 150 mM NaCl, 0.25% NP-40) containing 1 mM PMSF and 1X cOmplete® EDTA-free proteases inhibitors (Sigma) followed by centrifugation for 10 min at 15.800 g at 4 °C. Protein concentration was determined by the Bio-Rad Protein Assay (Bio-Rad) based on the Bradford method (Bradford 1976).

From 20 to 70 μg of total plant protein or from 10 to 20 μg of total purified recombinant protein was loaded per well in SDS-acrylamide/bisacrylamide gel electrophoresis using Tris-glycine-SDS buffer. Proteins were electrophoretically transferred to an Inmobilon-P polyvinylidene difluoride membrane (Millipore) using the Trans-Blot Turbo Transfer System (Bio-Rad). Membranes were blocked in Tris buffered saline-0.1% Tween 20 containing 5% blocking agent (semi-skimmed milk powder) and probed with antibodies diluted in blocking buffer with 1% blocking agent. For the detection of loading control, the membrane was incubated at 55°C for 10 min in stripping buffer (62.5 mM Tris-HCl pH 6.7, 2% SDS, 100 mM 2-mercaptoethanol) to remove previously bound antibody to re-probe it. Detection was performed using ECL Advance Western Blotting Detection Kit (Amersham) and the chemiluminescence was detected using an Intelligent Dark-Box II, LAS-1000 scanning system (Fujifilm). Alternatively, loading control was performed by staining the membrane with Ponceau S (Sigma-Aldrich) for 5 min before rinsing with water.

### *S*-nitrosylation assays

For *in vitro S*-nitrosylation, recombinant proteins were pretreated with the reducing agent DTT (20 mM) for 1 hr at room temperature in the dark to obtain the same redox status of cysteine residues in all the samples (-SH). Next, proteins were treated with the NO donors GSNO or/and SNAP (200 μM) for 30 min at room temperature in the dark with regular mixing. To check the reversibility of the modification, ultraviolet light was used as a reducing agent for 10 min under the same conditions after the GSNO/SNAP incubation. For - SH blocking, recombinant proteins were incubated with 20 mM *S*-methyl-methanethiosulfonate (MMTS) and 2.5% SDS at 50°C for 30 min with frequent mixing. Finally, 1 mM biotin HPDP (Pierce, Rockford, IL) and 1 mM ascorbic acid were added to the mixture and incubated for 1 hr at room temperature in the dark with regular mixing. After each step, reagents were removed by precipitation with 2 volumes of −20°C acetone following centrifugation at 2.500 g for 30 min at 4°C and the precipitated proteins were dissolved in 100 μL of HEN buffer (100 mM Hepes, 1 mM EDTA and 0.1 mM neocuproine, pH 7.8). Finally, recombinant proteins were loaded in SDS-acrylamide/bisacrylamide gel electrophoresis and transferred to a polyvinylidene difluoride membrane, to detect protein-SNO with anti-Biotin (Sigma, 1:4000).

For *semi-in vivo S*-nitrosylation of PAN, protein extracts from etiolated 7-day-old seedlings were obtained in extraction buffer (100 mM Tris-HCl, 150 mM NaCl, 0.25% NP-40) containing 1 mM PMSF and 1X cOmplete® EDTA-free proteases inhibitors (Sigma). Protein extracts (1 mg) were treated with SNAP (1 mM), GSNO (1 mM) or GSH (1 mM) for 1 hr in the dark and they were used as positive and negative controls, respectively. After, they were directly assayed by the biotin switch method. *In vitro* biotinylated proteins were purified by immunoprecipitation with 15 μL Protein A/G Ultralink resin (Pierce) per mg of protein with anti-biotin antibody for 2 hrs at 4 °C. The resin was pre-incubated with 2 μL of anti-biotin antibody (Sigma, 1:4,000). Beads were washed three times with HEN buffer (100 mM Hepes, 1 mM EDTA and 0.1 mM neocuproine, pH 7.8) and bound proteins were eluted with 10 mM DTT in SDS–PAGE solubilization buffer.

### *In vitro* denitrosylation assays

For denitrosylation assays, the resulting protein-SNO was incubated for 45 min with 5 μM TRX*h*5, 0.5 μM NTRA, and 1 mM NADPH as previously described by (Kneeshaw et al. 2014). Detection of different TGAs was performed by immunoblotting using an anti-Flag antibody. TRX*h*5 was detected with an anti-His tag antibody.

### Electrophoretic Mobility Shift Assay (EMSA)

EMSA studies using wild-type PAN and the cysteine-to-serine PAN variants *Cys340Ser*, *5 Cys to Ser* and *6 Cys to Ser* were performed as previously described in Gutsche et al., 2017. DNA-binding activity was analysed using a synthetic double-stranded oligonucleotide containing the palindromic TGA cis-regulatory element TGACGT flanked by a conserved GARP-binding, based on sequence features previously associated with PAN binding (Gutsche and Zachgo, 2016). The wild-type probe sequences were 5′-ATCAGATCTAATTGACGTGATAGCATATAGA-3′ and 5′-TCTATATGCTATCACGTCAATTAGATCTGAT-3′. A probe in which the TGACGT motif was replaced by TGATTT was used as a negative control (5′-ATCAGATCTAATTGATTTGATAGCATATAGA-3′ and 5′-TCTATATGCTATCAAATCAATTAGATCTGAT-3′). Oligonucleotides were 5′-labelled with 6-carboxyfluorescein (6-FAM).

For DNA-binding reactions, 1 µg of purified recombinant PAN protein was incubated with 3 ng pf the double-stranded DNA probe in binding buffer containing 100 mM Tris-KOH (pH 7.5), 500 mM KCl, 20% (v/v) glycerol and 0.05 µg µL⁻¹ poly(dI-dC) (Lindermayr et al., 2010) for 30 min at room temperature. For the redox EMSA studies and prior to the addition of the DNA probe, proteins were incubated with 1 mM GSNO (Sigma-Aldrich) for 30 min on ice. When analysing the removal of the –SNO modification, proteins were additionally incubated with 20 mM DTT for 20 min.

Free and protein-bound DNA were separated by electrophoresis on 6% polyacrylamide gels (40:1 acrylamide) prepared in 0.5× TBE and run at 4 °C. All EMSA experiments were independently repeated at least three times.

### Mass spectrometry

*In vitro* biotinylated proteins with SDS-PAGE solubilization buffer were loaded in 7% SDS-PAGE, visualised by Coomassie Blue Staining method and protein-stained bands were excised and manually digested with trypsin, chymotrypsin and endoproteinase AspN under non-reducing conditions as previously described(Albertos et al. 2015). After digestion, peptides were dried by speed-vacuum centrifugation and dissolved in loading solution (0.1% formic acid in water). Nano LC ESI-MSMS (liquid chromatography coupled to electrospray tandem mass spectrometry) analysis was performed using an Eksigent 1D-nanoHPLC coupled to a 5600 TripleTOF QTOF mass spectrometer (Sciex, Framinghan, MA, USA). The analytical column used was a silica-based reversed phase column Waters nanoACQUITY UPLC 75 μm I.D. × 15 cm, 1.7 μm particle size. The trap column was an AcclaimPepmap, 100 μm X 2 cm, 5 μm particle size, 100 Å pore size, switched on-line to the analytical column. The loading pump delivered a solution of 0.1% trifluoroacetic acid in 98% water/2% acetonitrile (Scharlab, Spain) at 2 μL/min. The nanopump provided a flow-rate of 250 nL/min and was operated under gradient elution conditions, using 0.1% formic acid (Fluka, Switzerland) in water as mobile phase A. and 0.1% formic acid in 100% acetonitrile as mobile phase B. Gradient elution was performed according to the following scheme: isocratic conditions of 96% A: 4% B for one minute, a linear increase to 50% B in 15 min, a linear increase to 90% B in 30 seconds, isocratic conditions of 90% B for five minutes and return to initial conditions in 30 seconds. Injection volume was 5 μL. The LC system was coupled via a nanospray source to the mass spectrometer. Automatic data-dependent acquisition using dynamic exclusion allowed to obtain both full scan (m/z 350-1250, 250msec) MS spectra followed by tandem MS CID spectra (100 msec) of the 15 most abundant ions per MS spectrum.

MS and MS/MS data were used to search against a customised target Arabidopsis protein database (31479 sequences) downloaded from UniprotKB. Database searches were done using a licensed version of Mascot v.2.5.1 and search parameters were set as follows: methylthio-cysteine, Biotin-HPDP (cysteine), Gln->pyro-Glu (N-term Q), Glu->pyro-Glu (N-term E), deamidation (NQ) and oxidised methionine as variable modifications; peptide mass tolerance was set at 25 ppm and 0.1 Da for MS and MS/MS spectra, respectively, and 2 missed cleavages were allowed. Mascot score threshold for peptide identification was set to a value equal or higher than 20. All MS/MS spectra corresponding to biotin-HPDP modified peptides were manually analysed.

### Tandem Affinity Purification assays (TAP)

Cloning of transgene encoding N-terminal GS^rhino^ tag (Van Leene et al. 2015) fusion under control of the constitutive cauliflower tobacco mosaic virus 35S promoter (*35S:GSrhino-GFP-PAN*) and transformation of Arabidopsis cell suspension cultures (PSB-D) with direct selection in liquid medium was carried out as previously described(Van Leene et al. 2011). TAP experiments were performed with 200 mg of total protein extract as input as described in Van Leene et al., 2015. Bound proteins were digested on bead after a final wash with 500 µL 50 mM NH_4_HCO_3_ (pH 8.0). Beads were incubated with 1 µg Trypsin/Lys-C in 50 µL 50 mM NH_4_OH and incubated at 37°C for 4 hrs in a thermomixer at 800 rpm. Next, the digest was separated from the beads, an extra 0.5 µg Trypsin/Lys-C was added, and the digest was further incubated overnight at 37°C. Finally, the digest was centrifuged at 20800 rcf in an Eppendorf centrifuge for 5 min, the supernatant was transferred to a new 1.5 mL Eppendorf tube, and the peptides were dried in a Speedvac and stored at −20°C until MS analysis. Co-purified proteins were identified by mass spectrometry using a Q Exactive mass spectrometer (ThermoFisher Scientific) using standard procedures. Proteins with at least two matched high confident peptides in at least 2 experiments were retained. Background proteins were filtered out based on frequency of occurrence of the co-purified proteins in a large dataset containing 543 TAP experiments using 115 different baits (Van Leene et al. 2015). True interactors that might have been filtered out because of their presence in the list of non-specific proteins were retained by means of semi-quantitative analysis using the average normalised spectral abundance factors (NSAF) of the identified proteins in the PAN TAPs (Van Leene et al. 2015).

### Statistical analysis

All statistical analyses were performed using R Statistical Software (v4.4.3; (R Core Team 2025). Outliers were identified using Rosner’s test implemented in the EnvStats package (Millard 2013). Statistical analyses and multiple-comparison procedures were performed using the car (Fox et al. 2024), rstatix (Kassambara 2023b), ARTool (Kay et al. 2021), emmeans (Lenth and Piaskowski 2026), multcomp (Hothorn et al. 2008), multcompView (Graves et al. 2024) and, nlme (Pinheiro et al. 2024) packages, as appropriate for each experimental design. Data visualisation was performed using ggplot2 (Wickham 2016) and ggpubr (Kassambara 2023a). The figures were exported using the officer and rvg (Gohel 2026) packages.

## Results

### PAN regulates root SCN in a NO-dependent manner

Using Arabidopsis as a model plant species, we aimed to decipher the effect of NO on PR growth and its impact on root architecture (Fernández-Marcos et al. 2011, 2012; Sanz et al. 2014). Considering the relevance of TFs in controlling root cell fate and architecture, we screened for TFs potentially involved in this process in the presence of a NO donor (SNAP) using the TRANSPLANTA collection of *β*-estradiol inducible lines (Coego et al. 2014). Further studies led to the identification of PERIANTHIA (PAN) as a potential regulator of root development (de Luis Balaguer et al. 2017; Hu et al. 2022).

First, we determined PR length in the *pan* mutant (SALK_051790), which has a loss-of-function mutation, and in the *XVE:PAN* transgenic line, which is inducible by *β*-estradiol, in the presence (SNAP) or absence (cPTIO) of NO. Although the genotype-treatment interaction was not significant, pairwise comparisons among treatments showed a significant PR reduction in the presence of SNAP in the case Col-0 and the *pan* mutant (Figure 1A). However, PAN activation resulted in dose-dependent inhibition of PR length in the presence of SNAP, which was partially restored in the presence of cPTIO (Figures 1A and S1A). These findings suggest that PAN may coordinate with other partners in regulating the function of the root SCN via NO.

**Figure 1.**
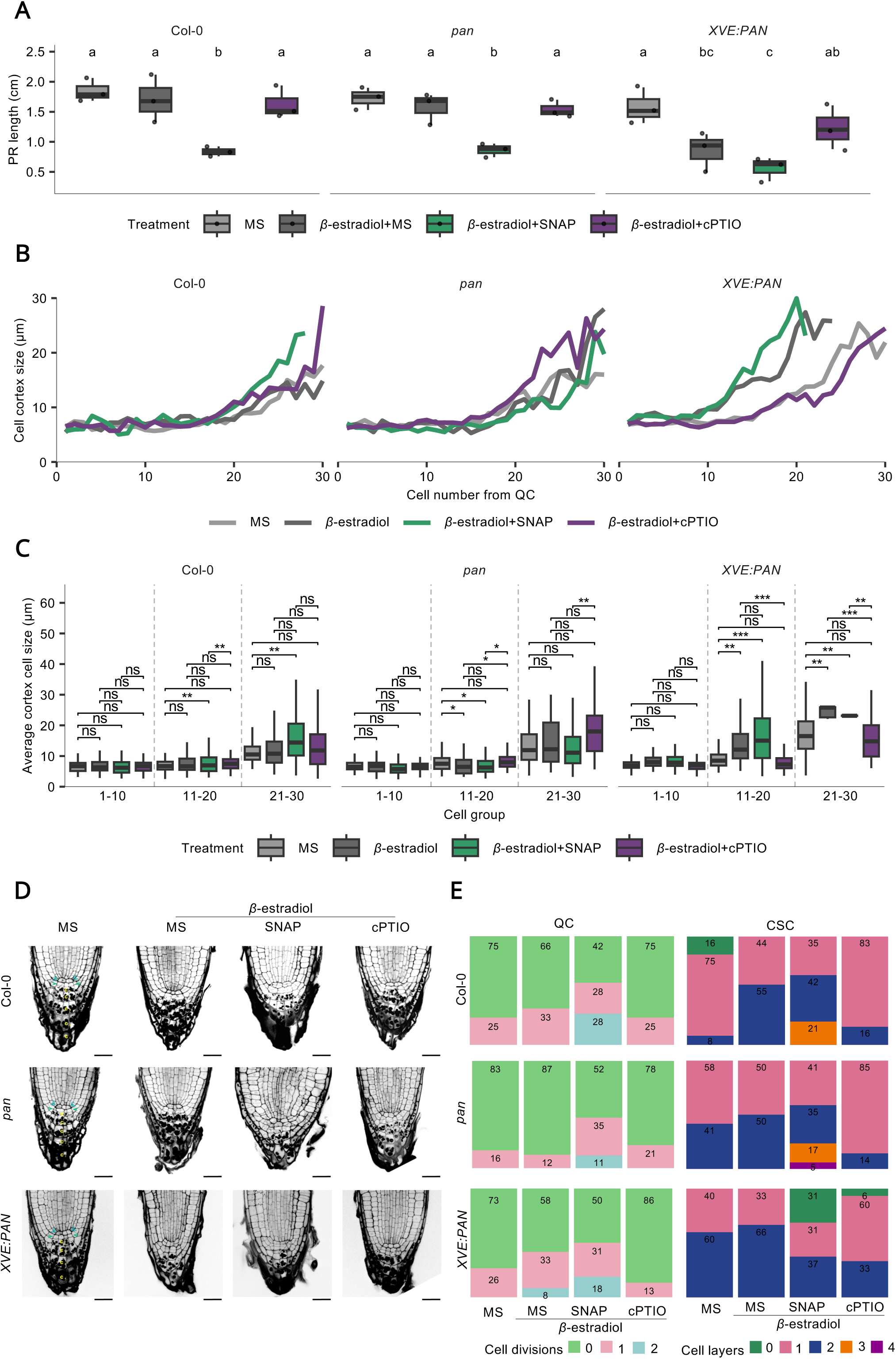
Nitric oxide regulation of RAM development involves PERIANTHIA (PAN). PR length and SCN analysis of Col-0, *pan* and *XVE:PAN* in 7-day-old seedlings grown on MS as control and MS supplemented with 5 µM *β*-estradiol and 5 µM *β*-estradiol in combination with 500 µM SNAP or 1 mM cPTIO. (A) Boxplot of PR length. Pairwise comparisons among treatments within each genotype were performed using estimated marginal means with Benjamini– Hochberg adjustment. Different letters indicate statistically significant differences (*P*<0.05). (N = 36). (B) Multiple line plots of the average cell size in the cortex cell layer (30 first cells from QC). (N≥3000). (C) Statistical differences were assessed using a three-way ART ANOVA, with root identity included as a random effect. Pairwise comparisons among treatments within each genotype and cell group were performed using estimated marginal means with Benjamini–Hochberg adjustment. Asterisks indicate statistically significant differences (*P* < 0.05). A total of 153 roots were analysed. (D) RAM confocal microscopy images. Roots were stained using the modified mPS-PI protocol. Blue triangles point to QC cells; green arrows point to CSC layers and yellow letters point to columella cell layers. Bars indicate 20 µm. (E) Stacked bar plot of the percentage of QC divisions and CSC layers. Different colours represent different numbers of QC divisions and/or cell layers. The numbers within the stacked bars correspond to the percentage. (N=144).

To uncover the cellular localization and expression pattern of PAN during early development, the *pPAN:GFP* transcriptional reporter line was used. The highest level of *PAN* transcripts was found in 2-day-old seedlings, with a decreasing trend from 5 days onwards (Figure S1B), although expression persisted at varying levels until flowering (Chuang et al. 1999). Building on our observation that PAN coordinates with NO to regulate root growth, we further investigated whether this gasotransmitter could affect the transcriptional profile or protein accumulation of PAN. Confocal imaging revealed that PAN was expressed exclusively in the QC (Figure S1C) with no statistically significant differences in *PAN* expression or localization under NO stimuli in short-term (6 hours) or long-term (24 hours) treatments (Figure S1D). We also examined whether short-term treatments (5 hours) with NO could transcriptionally modified the expression of *PAN*. No differences were found using whole plant tissue of *pPAN:GFP* transcriptional reporter line after of NO treatments (Figure S1E). Although our anti-PAN antibody was unable to detect endogenous PAN protein in roots due to its low abundance, the levels of transgenically expressed PAN protein were not affected by the addition or removal of NO (Figure S1F). These findings suggest that the expression of PAN in the QC is not directly modulated by NO and demonstrate that the putative molecular link between PAN and NO is independent of any effect on transcript and protein levels, thus pointing to a post-translational mechanism.

Given that root development depends on RAM activity, we explored whether the differences in phenotype were due to differences in the meristematic cell population. To this end, we used the elongation of meristematic cells as an indicator of cell differentiation (Pavelescu et al. 2018). Results showed that in the *XVE:PAN* inducible line (*β*-estradiol + MS), the elongation zone appeared around cells 11-13 (Figure 1B) and the largest cell size was in 11-20 and 21-30 of the cortex cells (Figure 1C). However, in *XVE:PAN*, the removal of NO (*β*-estradiol + cPTIO) led to restoration of cell cortex size values like Col-0 and the *pan* mutant (Figures 1B and 1C). In contrast, addition of NO (*β*-estradiol + SNAP) resulted in a significant increase in cell cortex size in *XVE:PAN* compared to Col-0 and the *pan* mutant (Figures 1B and 1C). These findings highlight that PAN promotes the differentiation of RAM cells in an NO dependent manner.

A detailed view of the SCN showed an increase in QC cell divisions in *XVE:PAN* line (*β*-estradiol + MS) compared to Col-0; while the *pan* mutant behaved slightly opposite. After NO treatment, Col-0 exhibited an increase of cell divisions in the QC, and of cell layers in CSC, confirming the role of NO in the root SCN. This behaviour slightly increased in the *pan* mutant and was partially restored in the case of *XVE:PAN* after NO addition (Figures 1D and 1E); whilst this effect was largely mitigated after NO scavenging (Figures 1D and 1E). Taken together, these results suggest that PAN has a significant effect over QC divisions and thus affects the subsequent differentiation of adjacent initial cells, namely columella cells, through a NO-dependent mechanism, resulting in an alteration of the primary root length.

### PAN is reversibly *S-*nitrosylated at multiple Cys residues

The identification of NO targets is crucial for understanding cellular redox regulation and the physiological role of this gasotransmitter. Interestingly, PAN protein contains six cysteines along its sequence, one of them (Cys340) located in the DOG1 C-terminal domain and the five remaining cysteine residues distributed throughout the PAN structure except in the bZIP domain (Figures S2A and S2B). Here, we demonstrate that recombinant PAN protein undergoes this PTM *in vitro* using the NO donor GSNO and the biotin switch technique (BST) (Figure 2A). Because Cys340 is an amino acid residue crucial for the function of PAN protein (Li et al. 2009), serine substitution of Cys340 (*Cys340Ser*) showed that this mutation did not affect to PAN nuclear location (Figure S2C) and that other cysteine residues may be key in the *S*-nitrosylation of PAN (PAN-SNO) (Figure 2B).

**Figure 2.**
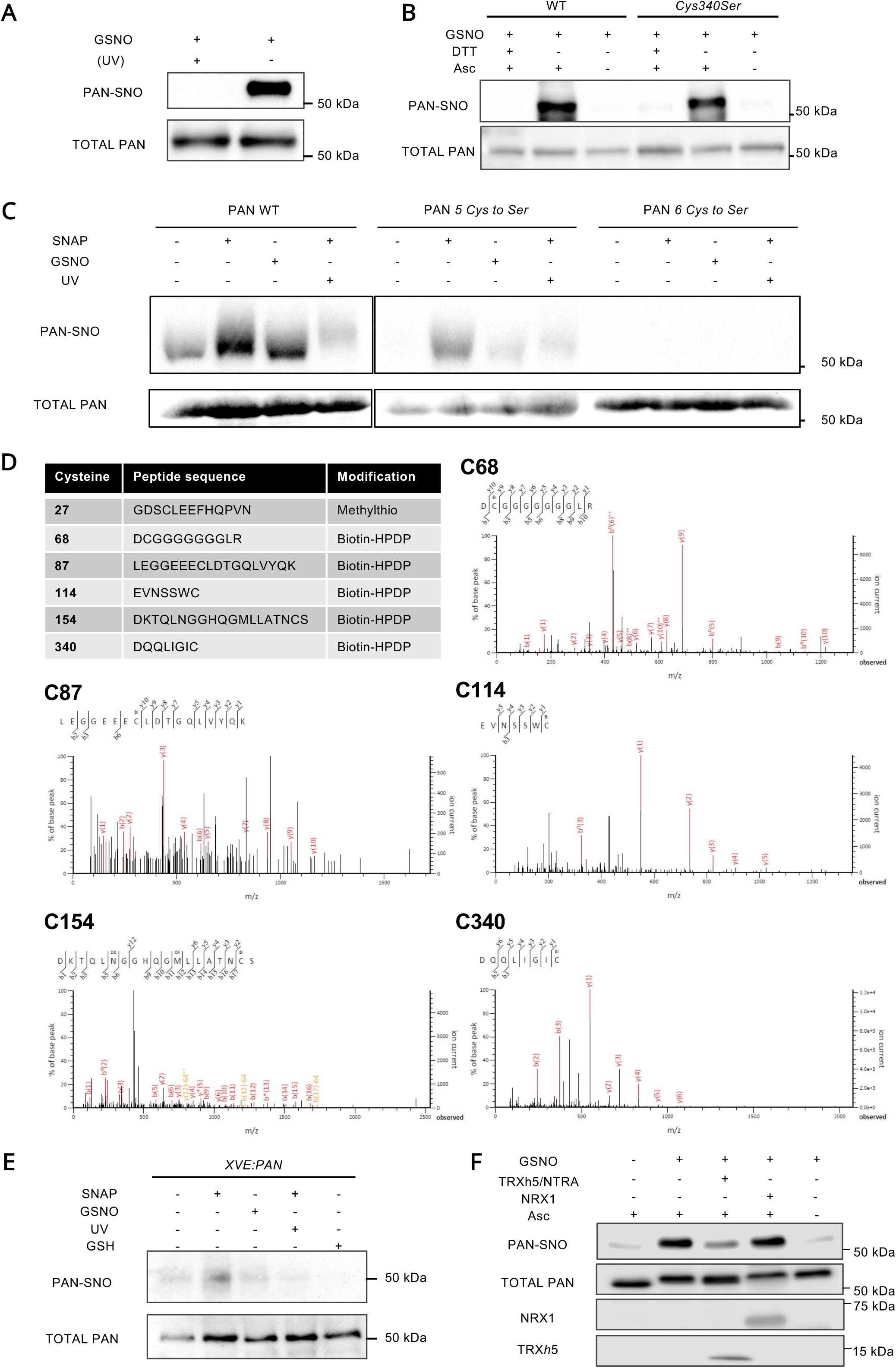
PAN is a target of *S*-nitrosylation. (A) *In vitro S*-nitrosylation of wild type (WT) PAN recombinant protein (PAN-SNO) using GSNO. (B) *In vitro S*-nitrosylation of WT and *Cys340Ser* PAN recombinant proteins using GSNO. (C) *In vitro S*-nitrosylation of WT, *5 Cys to Ser* and *6 Cys to Ser* PAN recombinant proteins by SNAP and GSNO using the biotin switch technique (BST). (D) Mass spectrometry analysis (LC-MS/MS liquid chromatography-mass spectrometry) of PAN-SNO cysteine modifications. Ion spectra of Cys 68, 87, 114, 154 and 340 modified by Biotin-HPDP. (E) Semi-*in vivo S*-nitrosylation of PAN using protein extracts from 7-day-old etiolated *XVE:PAN* seedlings incubated with SNAP or GSNO and analysed by BST. (F) Wheat-germ produced PAN protein was *S*-nitrosylated with GSNO and incubated with a combination of recombinant TRX*h*5, NTRA and NADPH. In all BST assays, SNAP and GSNO donors were used at 0.2 mM. Ultraviolet (UV) light was used as a SNO-reducing agent; SNO cleavage induced by 20 mM DTT and removal of sodium ascorbate (Asc) served as negative controls. Total PAN protein ensures equal protein loading.

To investigate the involvement of additional cysteine residues in PAN-SNO, two mutant versions were generated in which five (*C27C68C87C114C154*, PAN *5 Cys to Ser*) or six (*C27C68C87C114C154C340*, PAN 6 *Cys to Ser*) cysteines were replaced by serine amino acid. These variants were subsequently assessed for susceptibility to this NO-PTM. While the WT, *Cys340Ser* and *5 Cys to Ser* versions of PAN protein were *S*-nitrosylated, absence of SNO-signal was detected in the variant with no cysteines (PAN *6 Cys to Ser)* in the protein structure (Figure 2C). LC-MS/MS analysis confirmed that five of the six cysteines were *S*-nitrosylated (Figure 2D and Supplementary Dataset). Finally, *S*-nitrosylation was tested *in planta* using protein extracts from the *XVE:PAN* transgenic line treated with NO, thus demonstrating that plant-expressed PAN also harbours cysteine residues available to undergo this PTM (Figure 2E).

Based on the already documented reversibility of *S*-nitrosylation in plants by members of the thioredoxin (TRX) family (Kneeshaw et al. 2014), we further tested whether PAN *S*-nitrosylation was indeed reversible. In particular, the prototypical thioredoxin TRX*h*5 was previously reported to act as a protein-SNO reductase (Kneeshaw et al. 2014) and its expression is highly upregulated in the root cap, predominantly in columella cells (Belin et al. 2015). Considering the presence of PAN in QC cells and the importance of controlling the redox status of this key TF in the root SCN, we examined whether TRX*h*5 exhibits protein denitrosylation activity on PAN. Thus, wheat germ-produced PAN protein was *S*-nitrosylated by GSNO, desalted and incubated with a combination of recombinant TRX*h*5, recombinant NTR isoform A (NTRA) and NADPH (Kneeshaw et al. 2014) and the level of PAN-SNO was examined. Results showed that the fully reconstituted TRX*h*5/NTRA system was able to denitrosylate PAN *in vitro* (Figure 2F). These data indicate that *S*-nitrosylation of PAN is reversible and that protein-SNO reductase activity could potentially modulate PAN-SNO levels, thereby influencing CSC organisation and QC cell division in roots.

The TGA family of TFs regulate target gene expression by binding to specific *cis* regulatory elements containing the core sequence TGACGT, as previously demonstrated for PAN (Gutsche and Zachgo 2016). The effect of NO in the binding of PAN to this promoter sequence was assessed using Electrophoretic Mobility Shift Assays (EMSA) under reducing conditions (Figure S2D). PAN binding to the wild-type (WT) sequence was reduced following treatment with GSNO, whilst it was restored upon the addition of the reducing agent dithiotreitol (DTT). Following the *Cys340Ser* mutation, weak binding was observed in the presence of GSNO, demonstrating the inability of PAN to bind to the promoter in a NO-containing environment. By contrast, binding was clearly observed in the N-terminal 5 Cys mutated version (*5 Cys to Ser*), and similar binding was observed in the absence of cysteine residues in the amino acid sequence (*6 Cys to Ser*) (Figure S2D). As shown in Figure S2E, PAN binding to the TGACGT DNA motif was strongly dependent on the DTT concentration in the binding buffer while binding to the mutated version TGATTT (M) was completely abolished. Interestingly, the 5 Cys mutated version (only C340 present) and the *6 Cys to Ser* PAN protein without any cysteines showed a slightly different migration pattern compared to the WT and *Cys340Ser* protein indicating changes in folding for these variants. These results indicate that these Cys residues influence GSNO-mediated changes in the binding of PAN to DNA and suggest that *S*-nitrosylation affects the ability of PAN to bind to its targets.

### PAN differentially expressed genes co-regulate with NO scavenging

To investigate how NO regulates PAN activity during SCN development and root growth, transcriptional profiling of the root SCN was performed using RNA Sequencing (RNA-seq). The inducible *XVE:PAN* transgenic line was exposed to NO stimuli (the NO donor SNAP and the scavenger cPTIO) for 6 and 24 hours. Following quality control with FASTQC (Andrews, 2010), low-quality reads were removed, and the cleaned reads were then mapped to the Arabidopsis genome (TAIR10), achieving a mapping rate of >85% per sample. Differentially expressed genes (DEGs) were identified using a threshold of log2 fold change and q-value (logFC ≥ |1.5| and q-value ≤ 0.05). Regarding the number of DEGs, PAN induction by *β*-estradiol treatment alone resulted in 322 up-regulated and 367 down-regulated genes at 6h, whereas this response increased markedly when PAN induction was combined with cPTIO, reaching 552 up-regulated and 1,141 down-regulated genes. At 24h, *β*-estradiol treatment resulted in 593 up-regulated and 576 down-regulated genes, while the *β*-estradiol + cPTIO treatment further increased the transcriptional response to 702 up-regulated and 1,084 down-regulated genes (Figures 3A and S3A). Conversely, PAN induction in combination with the NO donor SNAP (*β*-estradiol + SNAP) elicited a limited transcriptional response at both time points (Figures 3A and S3A). Furthermore, volcano plots for the individual NO treatments with SNAP and cPTIO induced substantially weaker transcriptional changes than treatments involving PAN induction (Figure S3B). Taken together, these results suggest that NO availability modulates the transcriptome associated with PAN induction, with NO depletion causing the greatest transcriptional response.

**Figure 3.**
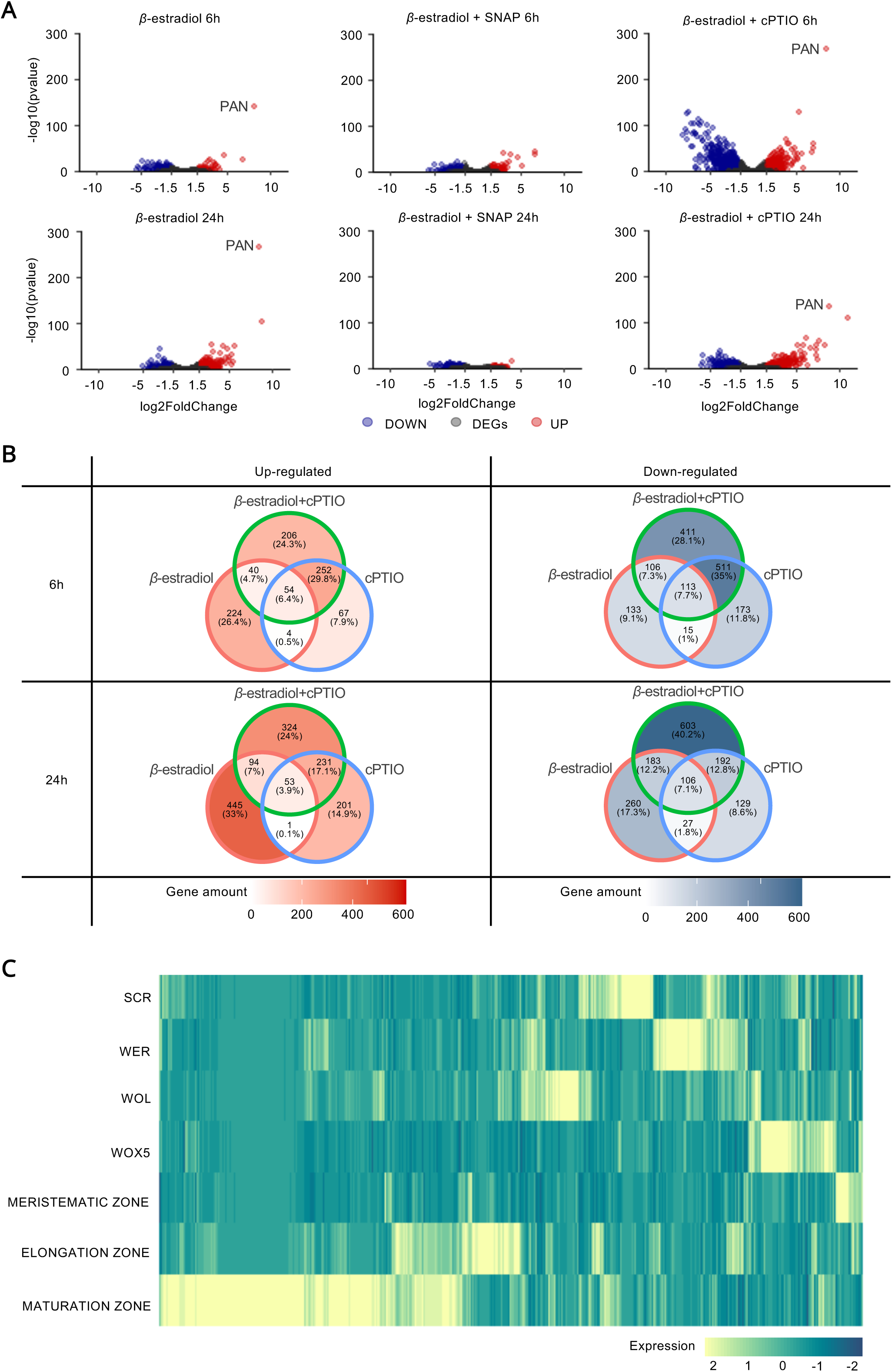
PAN transcriptional profile is co-regulated with NO depletion. (A) Volcano plots of significant DEGs under 5 µM *β*-estradiol, 5 µM *β*-estradiol in combination with 0.5 mM SNAP and 1 mM cPTIO for 6 and 24 hours. Red dots depict up-regulated genes (*qval < 0.05* and FC > 1.5), blue dots show down-regulated genes (*qval < 0.05* and FC < −1.5) and non-DEGs (*qval > 0.05)* are shown in grey. *PAN* is highlighted in all plots. (B) Venn diagrams indicating the percentage of common DEGs regulated by NO treatments. The 6- and 24-hour DEGs are displayed separately. Up-regulated DEGs (*qval < 0.05* and FC > 1.5) are highlighted in red, while down-regulated DEGs (*qval < 0.05* and FC < −1.5) are shown in blue. Colour intensity correlates with gene quantity. (C) Heatmaps of the locus identified in the 6-hour *β*-estradiol cPTIO treatments. Representation of normalised expression values (FPKM) obtained from the cell type transcriptome (Li et al. 2016).

Venn diagrams were used to illustrate the specific gene sets regulated by the different treatments. Notably 24.3% and 24% of the up-regulated genes at 6 and 24 hours, were uniquely induced by the combined application of *β*-estradiol and cPTIO and did not show any response to the individual treatments (Figure 3B). This particular response was even more evident in the set of down-regulated genes, where 28.1% (6 h) and 40.2% (24 h) of the total number of repressed genes were associated exclusively with the co-treatment (Figure 3B). Consistent with this, scatter plots confirmed an overall positive correlation among the treatments, as most genes exhibit the same direction of regulation (Figure S3C). However, the combination of *β*-estradiol and cPTIO modifies the transcriptional landscape by inducing or repressing a relevant subset of genes that are otherwise unresponsive to individual treatments. Since genes with opposing responses were in the minority, it appears that cPTIO modulates - rather than completely reverses-the transcriptional program associated with *β*-estradiol. In contrast, scarcely 1% of DEGs were common to all SNAP treatments, indicating a treatment-specific gene response (Figure S3D).

To identify development processes regulated by PAN in the absence of NO, the profiles of DEGs (*β*-estradiol + cPTIO) were compared with a previously published root expression map (Li et al. 2016). Interestingly, the greatest gene enrichment was observed in the root maturation zone (Figure 3C), suggesting that PAN promotes cell differentiation. Taken together, these data indicate that the transcriptional regulatory capacity of PAN is modulated by the availability of NO in the root SCN.

### TGA members interact with PAN, are *S*-nitrosylated differentially and contribute to the NO sensing in the root SCN

PAN is a member of the TGA TF family, as previously reported (Chuang et al. 1999; Maier et al. 2011). Evidence suggests that the plant bZIP family can form homo- and heterodimers, providing regulatory flexibility in target site selection and protein interactions (Lorenzo 2019; Li et al. 2023). To determine which proteins interact with PAN *in vivo*, protein complexes were first analysed via tandem affinity purification (TAP) (Van Leene et al. 2015) and then identified by mass spectrometry (Supplementary Table 1). The results showed proteins belonging primarily to the TGA family; specifically, TGA2, TGA3 and TGA5 were detected in both replicates, while TGA1, TGA6, and TGA7 were found only in the second analysis. To confirm the findings obtained by TAP-tagging analysis, these interactions were validated *in planta* using Bimolecular Fluorescence Complementation (BiFC) assays. The results demonstrated the interaction of PAN with TGA2, TGA3 and TGA5, as well as the nuclear localization of these complexes in tobacco leaf cells (Figure S4A). Consequently, we conclude that other TGA family members, specifically TGA2, TGA3 and TGA5, function as PAN interactors.

Since *S*-nitrosylation of TGA1 has been previously described (Lindermayr et al. 2010) and *S*-nitrosylation of PAN has been reported in this study, we investigated whether other TGAs are targets of this redox-based PTM. To determine the susceptibility of TGA1, TGA2, TGA3, TGA5, TGA6, TGA7 and PAN to *S*-nitrosylation, wheat germ-produced proteins were exposed to GSNO and the formation of TGA-SNO was monitored using the BST method. As shown in Figure 4A, TGA1, TGA2, TGA3, and TGA7 were sensitive to *S*-nitrosylation, whereas TGA5 and TGA6 were not. The *S*-nitrosylation of PAN was confirmed under identical conditions, indicating that multiple members of the TGA family are susceptible to this posttranslational modification.

**Figure 4.**
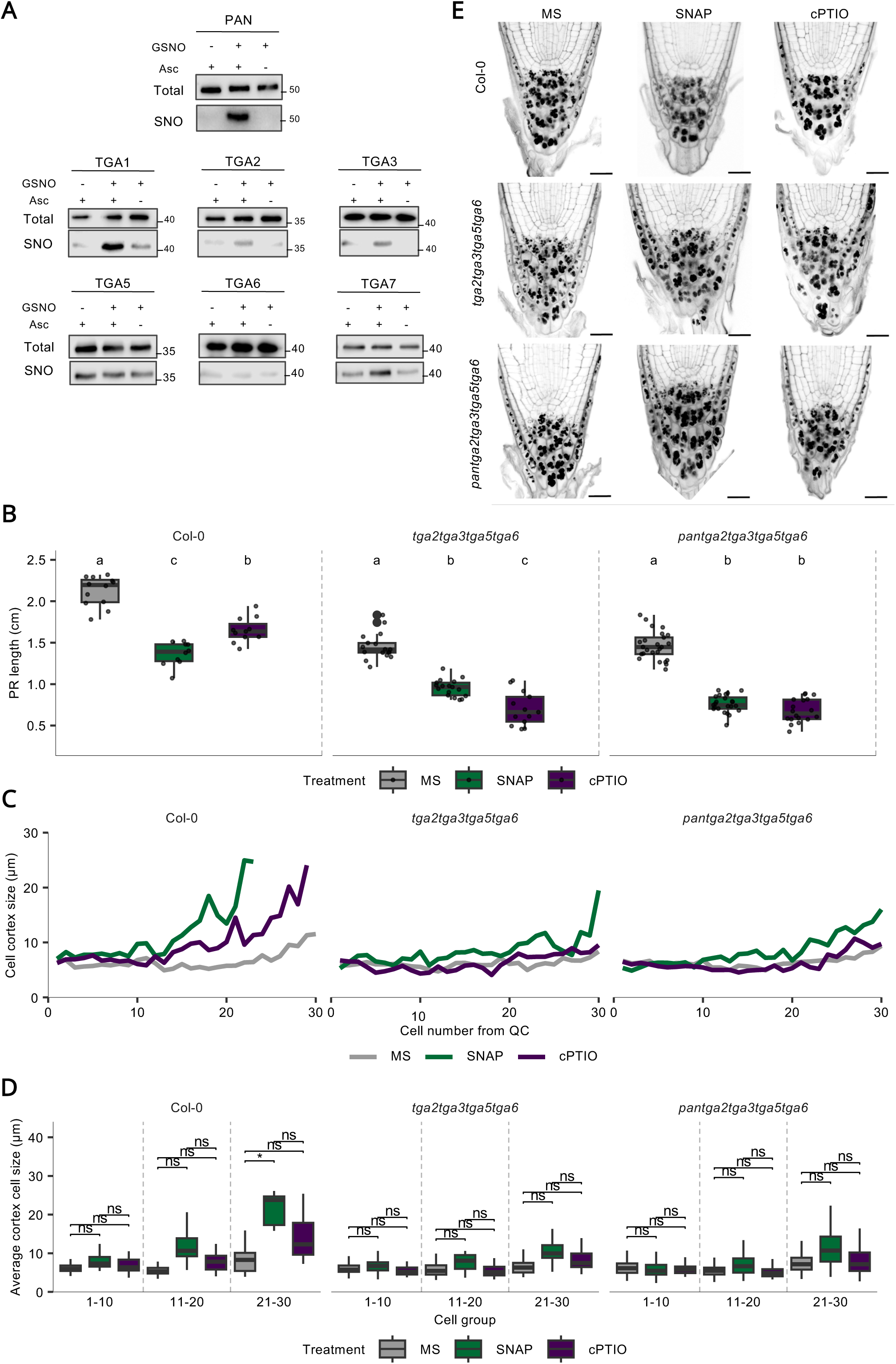
Members of the TGA family participate in NO sensing during RAM development. (A) *In vitro S*-nitrosylation of PAN interactors from the TGA family. Wheat-germ produced proteins were incubated with 1 mM GSNO, and SNO modification was detected by BST. Sodium ascorbate (Asc) was used to specifically detect *S*-nitrosylated proteins, and the absence of this reducing agent was used as a negative control. TGA proteins were detected using the α-His antibody. Total represents input loading control, and SNO shows the biotin/SNO-captured protein. (B) Boxplot of PR length of 7-day-old Col-0, *quadruple* and *pentuple* seedlings grown on MS, SNAP or cPTIO. Pairwise comparisons among treatments within each genotype were performed using estimated marginal means with Benjamini–Hochberg adjustment. Different letters indicate statistically significant differences (*P*<0.05). (N = 132). (C) Multiple line plots of the average cell size in the cortex cell layer (first 30 QC cells) of 7-day-old seedlings roots of Col-0, *quadruple* and *pentuple* mutants grown on MS, 0.5 mM SNAP or 1 mM cPTIO. (N≥2000). (D) Bar plots showing the average size of cortex cells grouped according to their position from QC in 1-10, 11-20 and 21-30 of roots from 7-day-old seedlings of Col-0, *quadruple* and *pentuple* mutants grown on MS, 0.5 mM SNAP or 1 mM cPTIO. Statistical differences were assessed using a three-way ART ANOVA, with root identity included as a random effect. Pairwise comparisons among treatments within each genotype and cell group were performed using estimated marginal means with Benjamini–Hochberg adjustment. Asterisks indicate statistically significant differences (*P* < 0.05). A total of 56 roots were analysed. (E) RAM confocal microscopy images of SCN from 7-day-old roots of Col-0, *quadruple* and *pentuple tga* mutants under control conditions or treated with SNAP or cPTIO. Roots were stained using the modified mPS-PI protocol.

The spatial expression pattern of PAN in Arabidopsis root QC cells prompted an investigation into whether other members of the TGA family exhibited a similar transcriptional profile. Although TGA-type TFs have been shown to be involved in root development (Hu et al. 2022), their regulatory role in the maintenance and spatial organization of the root SCN remains unknown. Data from the available transcriptomic profile of root tissues (de Luis Balaguer et al. 2017) revealed that other TGA members are expressed, at varying levels, in root stem cells (Supplementary Table 2). PAN expression is enriched 8-fold in the SCN and 4-fold in QC cells compared to the elongation and differentiation zones, respectively (de Luis Balaguer et al. 2017). However, most TGAs are expressed but not enriched in SCN-related cells, except for TGA7, which showed no differential expression compared to the differentiation zone (Supplementary Table 2). Therefore, to examine the contribution of PAN and other TGA members to the NO-mediated regulation of root meristem organisation by NO, PR growth was analysed in 7-day-old *tga*-related mutants. The *tga7* mutant exhibited similar PR growth to Col-0 under control and NO treatments; however, root growth was severely impaired by the cumulative mutations in other members of the TGA family, both under control conditions and after pharmacological treatment with NO (Figure S4B). The reduction in PR length was particularly evident following NO removal, where both the *triple* (*tga2tga3tga5*) and *quadruple* (*tga2tga3tga5tga6*) *tga* mutants demonstrate the necessity of this family of TFs for NO sensing. To determine whether the *pan* mutation contributes to this reduction in PR growth, a *pan* mutant was crossed with a *quadruple tga* mutant to obtain the *pantga2tga3tga5tga6* (*pentuple*) mutant (Figure S4C). It is worth noting that mutations in several TGA members negatively affected the development of adult Arabidopsis plants, resulting in smaller rosettes and shorter stems in *quadruple* and *pentuple* mutants (Figure S4D). A detailed analysis revealed a significant genotype-treatment interaction, indicating that the effect of treatment on PR growth varied depending on the genotype. Indeed, Hu et al. (2022) reported that *quadruple tga* mutants exhibited a significant reduction in PR growth, which was corroborated under our experimental conditions (Figure 4B). Interestingly, analysis of the *pentuple tga* mutant revealed further shortening of the PR length, which decreased drastically following the addition and scavenging of NO (Figure 4B), highlighting the necessity of the TGA family of TFs in the sensing of this gasotransmitter for proper RAM development. Furthermore, a significant genotype-treatment-cell group interaction was detected for cortex cell size (*P* < 0.001), suggesting that the impact of treatment varies according to genotype and cell position. Therefore, the effects of treatment were analysed separately within each genotype and cell group. The overall results regarding cortex cell size showed that the *pentuple tga* mutant was insensitive to both addition and depletion of NO (Figures 4C and 4D), which further emphasises the importance of the TGA family of TFs in regulating root growth via NO. Beyond primary growth, we examined whether the structural defects in the root meristem of *tga* mutants affected root functionality. Consistent with the disorganisation observed in the SCN, analysis of the gravitropic response revealed an additive effect on PR curvature as TGA mutations were introduced (Figure S4E), further supporting functional cooperation within this family of TFs. Due to the distortion of the SCN pattern observed in the *quadruple* and *pentuple tga* mutants, it was not possible to accurately quantify the cell divisions happening in the QC and CSC. Nevertheless, an anomalous organisation of these stem cells was observed, along with premature differentiation of the CSC. The addition and removal of NO had no additional effect on SCN organisation compared to untreated *quadruple* and *pentuple tga* mutants (Figure 4E), these results being consistent with those described above. Overall, the data presented here demonstrate that transcription factors of the TGA family, predominantly orchestrated by PAN activity, are necessary for the sensing and integration of NO-mediated sensing and signalling, thus leading to the proper organization of the root SCN.

**Figure 5.**
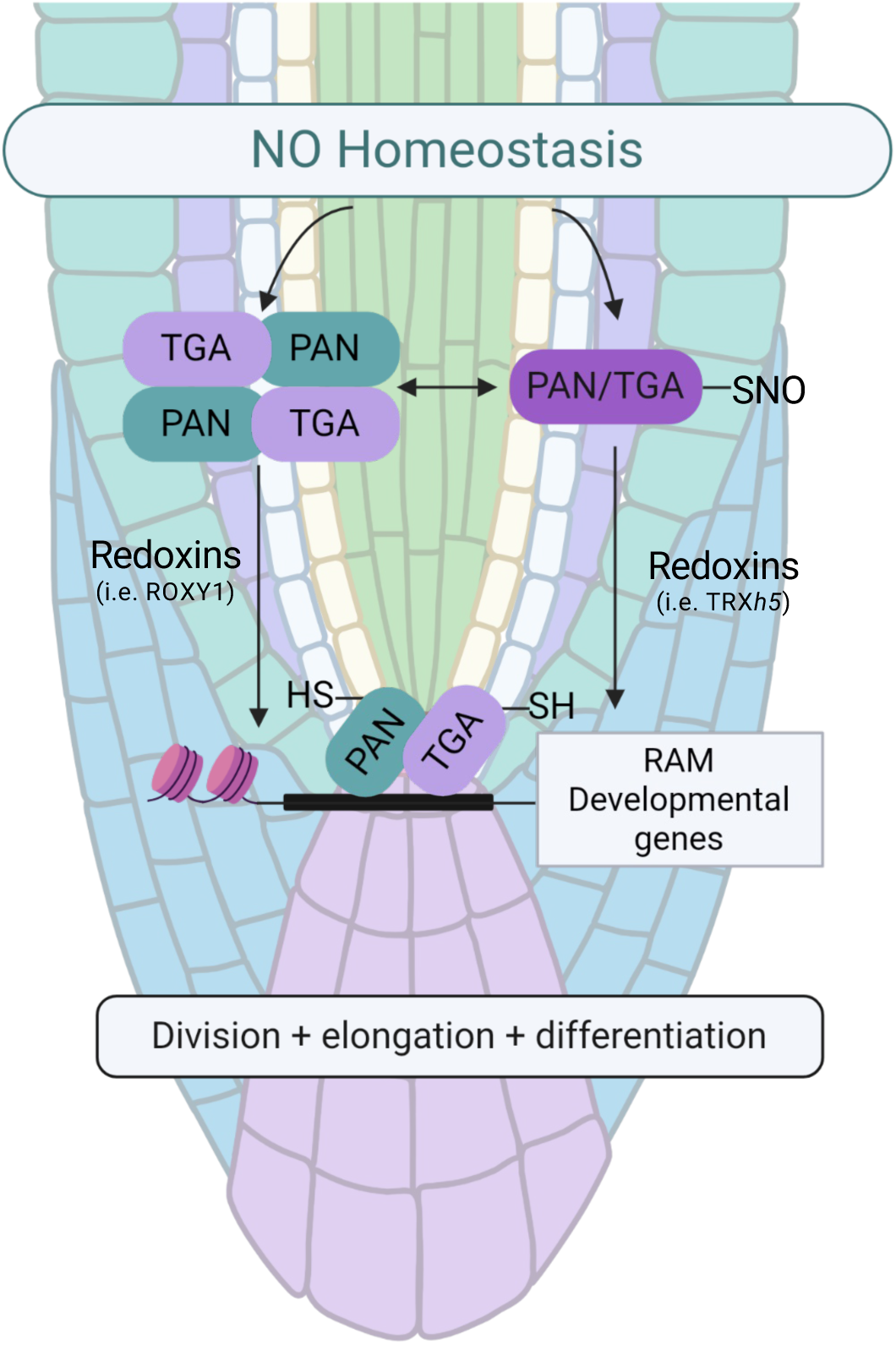
Model showing the role of NO in the PAN-based regulatory mechanism during RAM development. The maintenance of RAM in plants depends, although not exclusively, on NO homeostasis. We have previously reported that NO affects cell meristem size and number (Fernández-Marcos et al. 2011, 2012, 2013) and plays an important role in regulating stem cell decisions (Sanz et al. 2014). Furthermore, our previous studies have identified PAN as a molecular hub in the root SCN being involved in QC function and CSC maintenance(de Luis Balaguer et al. 2017). Here we show that *S*-nitrosylation of PAN (and other TGA transcription factors) inhibits its binding capacity to DNA and associated transcriptional activation. Independently, other redox signals can modify PAN structure (i.e. ROXY1 (Gutsche and Zachgo 2016)). Thus, both the oligomeric form of PAN and *S*-nitrosylated PAN (PAN-SNO) are established as reservoirs of this transcription factor. With the appropriate developmental signal and the action of specific reductase enzymes (i.e. TRX*h*5), PAN, together with other TGA partners, can bind to promoters containing the TGA motif and thus modify the gene expression responsible for maintaining RAM, including the homeodomain transcription factor necessary for the function of the quiescent centre (QC).

## Discussion

PAN has been established as a central hub in the organisation and development of SAM (Maier et al. 2009, 2011) and, more recently, as an essential component of the regulatory network governing RAM development (de Luis Balaguer et al. 2017). While PAN transcription is limited to the QC (Nawy et al. 2005; Lee et al. 2006), previous data indicate that the QC is an NO-free environment (Fernández-Marcos et al. 2011; Sanz et al. 2014), contrasting with the accumulation of NO in the cortex and endodermis of stem cells and their immediate progeny, generating endodermal and cortical tissues (Sanz et al. 2014). This spatial distribution suggests that the QC acts as a protective niche for PAN activity, where the absence of NO prevents its early inhibition. Conversely, the presence of NO in the surrounding tissues could function as a post-translational “off switch,” limiting the influence of PAN on the SCN and promoting differentiation as cells move away from the QC.

Our results indicate that the activation of PAN inhibits PR length, a phenotype partially reversed by NO scavenging (cPTIO) but not by the addition of NO (SNAP). Given that RAM activity and the elongation of meristematic cells are considered an indicator of cell differentiation (Pavelescu et al. 2018), we investigated whether the aforementioned phenotype was due to differences in the meristematic cell population. The results shown here indicate that cell cortex size returns to the levels observed in Col-0 and the *pan* mutant only when NO is removed. This suggests that PAN promotes the differentiation of RAM cells by regulating QC divisions and, consequently, influences the subsequent differentiation of adjacent cells through a NO-dependent mechanism. We agree that ectopic expression of PAN may occur when using the *XVE:PAN* inducible line. However, the differential response observed between *β*-estradiol alone and its combination with cPTIO justifies this approach. Whilst *β*-estradiol itself induces only a mild phenotypic and transcriptomic response, the differential behaviours between treatments and genotypes reinforce the specific role of PAN in the regulation of root SCN. Based on the functional link between PAN and NO, we analysed both the transcript and protein levels of PAN in QC cells and found that these remained steady regardless of NO treatment. Instead, the regulatory function of PAN seems to be modulated at the post-translational level, as PAN undergoes *S*-nitrosylation. It has been already described that this PTM modifies other members of the bZIP family via NO, including ABI5, where NO reduces ABI5 protein levels, thereby enabling seed germination (Albertos et al. 2015, 2021) or bZIP67, where *S*-nitrosylation stabilises this TF thus controlling the seed fatty acid profile (Sánchez-Vicente et al. 2024); as well as TGA1, whose DNA-binding activity is enhanced by *S*-nitrosylation (Lindermayr et al. 2010). Consequently, this NO-PTM alters the structure and activity of TFs, influencing their ability to bind to DNA and interact with other proteins, thereby setting the functional limits of their activity. It is worth noting the reversibility of this post-translational modification, which has been described in plant systems (Kneeshaw et al. 2014), as this adds another level of regulation. The activity of TRX*h*5, together with NADPH-dependent thioredoxin reductase A (NTRA), can reverse the *S*-nitrosylation of PAN. This reveals an NO-dependent ‘on-off’ mechanism that fine-tunes QC activity, consequently affecting the functioning of the RAM. However, it could be argued that, if *S*-nitrosylation of PAN were the sole critical factor for root growth, the *pan* mutant should show greater tolerance to SNAP and cPTIO than the wild type or the inducible lines. Nevertheless, the fact that this has not been observed is probably due to the low penetrance (10-12%) and the weak allelic nature of the *pan* mutation (de Luis Balaguer et al. 2017), together with functional synergy with other TGA members as described in this study. Our EMSA analyses also indicate that NO modulates the DNA-binding activity of PAN. GSNO was found to reduce PAN binding to a synthetic sequence containing the TGACGT motif, an effect that was reversed by DTT. This suggests that the inhibition is sensitive to the redox environment. The differential behaviour of PAN variants (*Cys to Ser* mutants) further suggests that specific *Cys* residues are involved in this regulation. These findings are consistent with a model in which *S*-nitrosylation modulates PAN binding to the regulatory sequences of its target genes, consequently affecting PAN-dependent transcriptional responses in the root SCN (Figure 6).

RNA sequencing data suggest that the availability of NO significantly influences the transcriptional response associated with PAN activation. Reduced NO availability increased the number of DEGs associated with PAN, while SNAP treatment attenuated this response. Alongside the results of the EMSA and *S*-nitrosylation assays, these findings imply that NO could regulate PAN activity precisely at the transcriptional level, potentially by altering its capacity to bind to target *cis*-regulatory elements. Moreover, nearly 25% of the DEGs in the root SCN were up-or down-regulated exclusively under this condition. These findings lend weight to the hypothesis that the regulatory capacity of PAN may generate a signal that inhibits its activity, most likely via *S*-nitrosylation, which is consistent with our transcriptomic data. In parallel, we identified a correlation between DEGs and the root tip structure through expression analyses along the root map (Li et al. 2016), with the maturation zone being the most significantly affected. These results corroborate the hypothesis proposed by de Luis Balaguer et al., 2017 where the expression of stem cell regulators gradually decreases from the SCN to the meristematic zone.

Given that bZIP heterodimerization is known to significantly alter DNA-binding specificity (Li et al. 2023), we have demonstrated that PAN interacts with other members of the TGA family, some of which are also susceptible to *S*-nitrosylation. To assess the biological significance of this interaction with other TGAs, the phenotypic effect of the cumulative mutation of various TGAs on PR length was investigated. Single *tga* mutants showed behaviour like that observed in Col-0 regarding PR length, whilst, as shown here and previously described, the single *pan* mutant exhibits a significant reduction in PR length (de Luis Balaguer et al. 2017; Hu et al. 2022) highlighting its relevance in the maintenance of the root SCN. Here, we confirmed a drastic reduction in PR growth as *tga* mutations accumulate in *quadruple* and *pentuple tga* mutants, as previously demonstrated by Hu et al., 2022, whilst the addition and removal of NO had no further effect on PR length or SCN organization compared with untreated *quadruple* and *pentuple tga* mutants. This redundancy suggests that PAN should not be regarded as an isolated “master hub,” but rather as an essential integrator within a regulatory node that depends on the TGA family and is sensitive to cellular redox status. This redundancy explains why the *pentuple* mutant is required to fully manifest the phenotypic consequences of these interactions. Overall, these findings suggest that other members of the TGA family are essential, alongside PAN, for the sensing of NO during root development. Taken together, these studies emphasize the role of NO in modulating root growth and the fate of root stem cells by regulating PAN/TGA activity (Figure 6).

## AUTHOR CONTRIBUTIONS

S.G.J. and M.G.F.-E. performed research; A.S-C. performed confocal images of root SCN. C.M.-P. and S.H.S. performed TGA-SNO biotin switch. N.G. and S.Z. performed EMSAs; J.V.L and G.J. performed TAP-tagging assays; S.G.J., J.S., M.A.L.B and R.S. carried out the RNAseq analysis. M.G.F.-E., S.G.J., C.M.-P. and O.L. analysed data; C.M.P. and O.L. designed research, supervised the work and wrote the paper and O.L. provided financial support. All authors discussed the results and commented on the manuscript.

## ACKNOWLEDGEMENT

We thank the Spanish networks RED2022-134072-T and RED2022-134917-T for critical comments and stimulating discussions of the manuscript. We also thank proteomics facility CNB-CSIC for technical proteomic assistance, VIB’s Proteomics Expertise Centre (PEC) for MS identification and Dominique Eeckhout for analysis of the MS experiments, and Sequencing Facility of NUCLEUS, University of Salamanca.

## Supplementary Material

**Supplementary Table 1.** Identification of GS^rhino^-PAN interacting proteins by TAP purification followed by tandem mass spectrometry (TAP-MS). E-1, NSAF value in experiment 1; E-2, NSAF value in experiment 2; T, total number a protein was found in the two TAP experiments.

| Gene | Description | E-1 | E-2 | T |
| --- | --- | --- | --- | --- |
| AT1G68640 | PAN | 0.577 | 1.202 | 2 |
| AT5G06950 | TGA2 | 0.425 | 0.604 | 2 |
| AT5G06960 | TGA5 | 0.394 | 0.569 | 2 |
| AT1G22070 | TGA3 | 0.048 | 0.080 | 2 |
| AT3G12250 | TGA6 |  | 0.538 | 1 |
| AT1G77920 | TGA7 |  | 0.056 | 1 |
| AT5G65210 | TGA1 |  | 0.055 | 1 |
| AT1G02840 | Serine arginine-rich splicing factor 34 (SRP34) | 0.062 |  | 1 |
| AT1G04600 | Myosin XI A (XIA) | 0.011 |  | 1 |
| AT1G16470 | Proteasome subunit (PAB1) | 0.081 |  | 1 |
| AT1G22840 | Cytochrome C-1 (CYTC-1) | 0.168 |  | 1 |
| AT1G23860 | Serine/arginine-rich-containing zinc finger protein 21 (RSZ21) | 0.097 |  | 1 |
| AT1G56340 | Calreticulin 1 (CRT1) | 0.107 |  | 1 |
| AT1G73230 | Nascent polypeptide-associated complex NAC | 0.289 |  | 1 |
| AT2G07727 | Cytochrome B/B6 | 0.047 |  | 1 |
| AT2G24200 | Leucyl aminopeptidase 1 (LAP1) | 0.038 |  | 1 |
| AT2G24590 | Serine/arginine-rich-containing zinc finger protein 22A (RSZ22A) | 0.142 |  | 1 |
| AT2G40660 | Nucleic acid-binding, OB-fold-like protein | 0.049 |  | 1 |
| AT2G44160 | Methylenetetrahydrofolate reductase 2 (MTHFR) | 0.078 |  | 1 |
| AT3G10920 | Manganese superoxide dismutase 1 (MSD1) | 0.163 |  | 1 |
| AT3G11710 | Lysyl-Trna synthetase 1 (ATKRS1) | 0.044 |  | 1 |
| AT3G23940 | Dihydroxyacid dehydratase (DHAD) | 0.032 |  | 1 |
| AT3G25230 | Rotamase FKBP 1 (ROF1) | 0.051 |  | 1 |
| AT3G27240 | Cytochrome C1 family | 0.062 |  | 1 |
| AT3G51800 | ERBB-3 binding protein 1 (EBP1) | 0.048 |  | 1 |
| AT3G52300 | ATP synthase D chain, mitochondrial | 0.159 |  | 1 |
| AT3G53460 | Chloroplast RNA-binding protein 29 (CP29) | 0.058 |  | 1 |
| AT3G53580 | Diaminopimelate epimerase | 0.053 |  | 1 |
| AT4G13430 | Isopropyl malate isomerase large subunit 1 (IIL1) | 0.038 |  | 1 |
| AT5G10540 | Thimet metalloendopeptidase 2 (TOP2) | 0.039 |  | 1 |
| AT5G12110 | Glutathione s-transferase, C-Terminal-like / Elongation factor EF1B | 0.084 |  | 1 |
| AT5G13520 | Peptidase M1 family protein | 0.045 |  | 1 |
| AT5G14590 | Isocitrate/isopropylmalate dehydrogenase | 0.058 |  | 1 |
| AT5G25450 | Cytochrome BD ubiquinol oxidase | 0.142 |  | 1 |
| AT5G37600 | Glutamine synthase (GLN1;1) | 0.053 |  | 1 |
| AT5G40810 | Cytochrome C1 family | 0.093 |  | 1 |
| AT5G47210 | Hyaluronan | 0.082 |  | 1 |
| AT5G53560 | Cytochrome B5 Isoform E (CB5-E) | 0.207 |  | 1 |
| AT5G61790 | Calnexin 1 (CNX1) | 0.052 |  | 1 |

**Supplementary Table 2.** Cell type-specific TGA transcriptional data from the QC, CEI, XYL, and the whole SCN compared to the transcriptional profiles corresponding to elongation and differentiation zones of the root. CEI: cortex/endodermis initial cells; XYL: xylem initial cells.

|  | SCN | CEI | XYL | QC | Elongatio<br>n | Differentiatio<br>n |
| --- | --- | --- | --- | --- | --- | --- |
| **TGA1** | 1.053 | 1.559 | 0.788 | 0.960 | 0.892 | 1.806 |
| **TGA3** | 1.182 | 0.893 | 0.921 | 0.848 | 1.044 | 1.872 |
| **TGA2** | 1.294 | 0.993 | 1.091 | 1.433 | 0.916 | 0.815 |
| **TGA5** | 1.720 | 1.131 | 1.238 | 1.493 | 1.597 | 1.537 |
| **TGA6** | 0.849 | 0.730 | 0.730 | 0.793 | 0.858 | 0.833 |
| **TGA7** | 2.993 | 0.870 | 0.508 | 1.556 | 1.298 | 3.089 |
| **PAN** | 4.138 | 1.113 | 0.560 | 2.790 | 0.551 | 0.529 |

**Supplementary Table 3.**
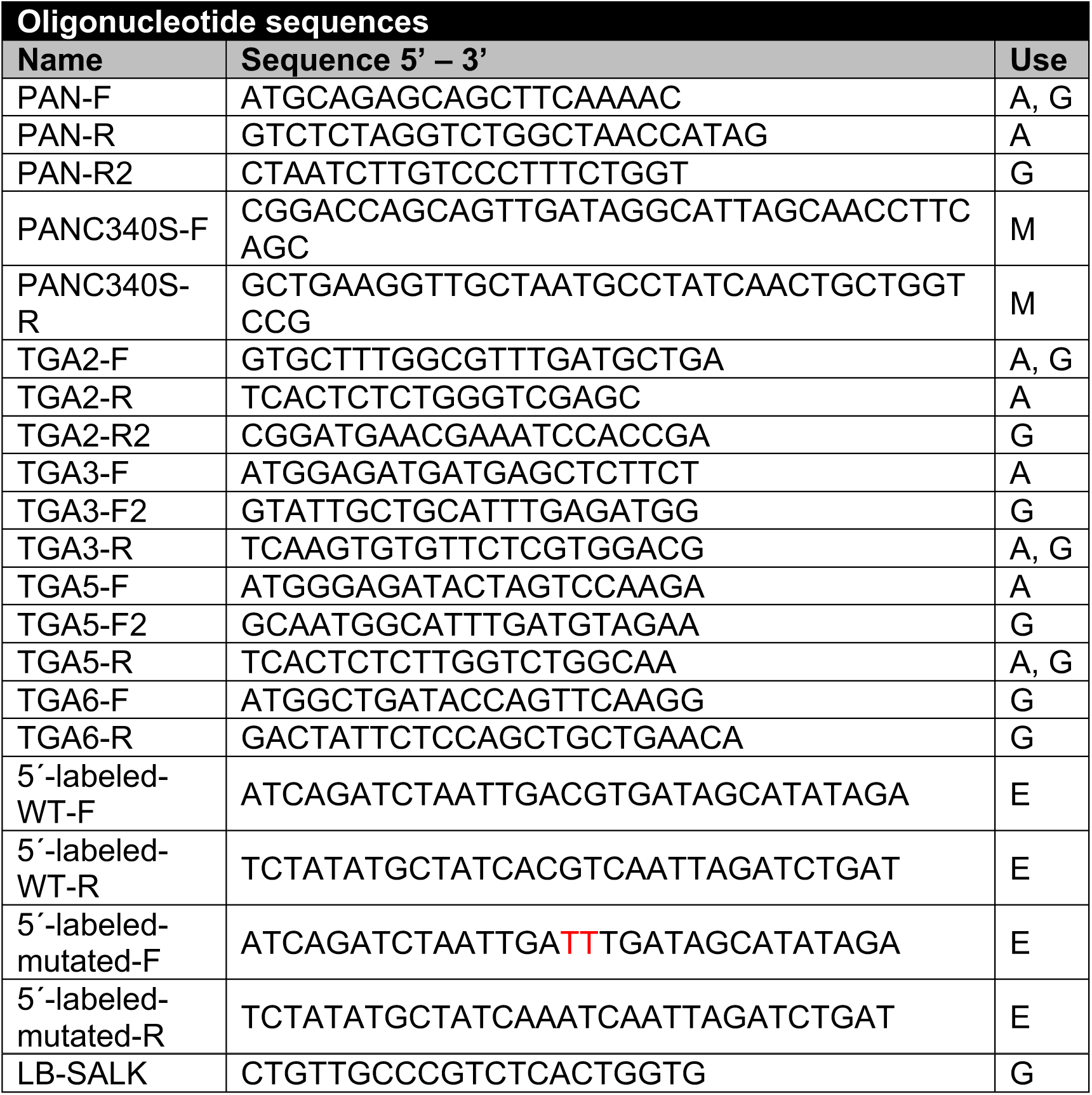
List of oligonucleotides used as PCR primers. F, forward; R, reverse; A, amplify CDS; G, genotyping; M, site-directed mutagenesis; EMSA, E.

## Data availability statement

Transcriptome sequence reads have been deposited in the GEO database under BioProject number GSE212583 (TGA family of transcription factors are nitric oxide sensors in the root stem cell niche).

## Supplementary Figure Legends

**Supplementary Figure 1 (S1).**
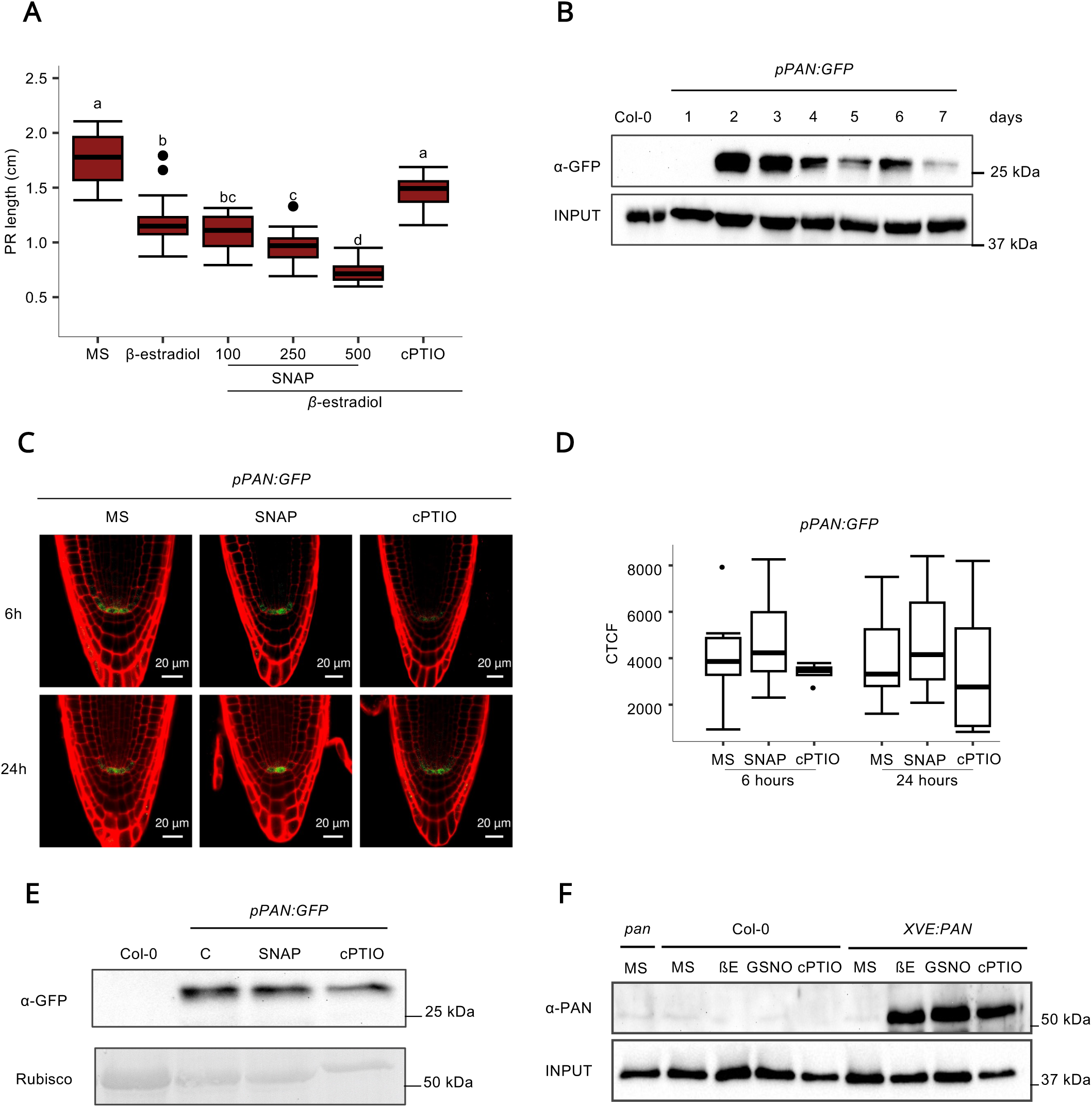
Effect of NO on transcription and protein levels of *PAN* in QC cells. *Supporting Figure 1*. (A) Boxplot of the PR length of 7-day-old *XVE:PAN* seedlings grown in MS as control and MS supplemented with 5 µM *β*-estradiol or 5 µM *β*-estradiol in combination with different concentrations of SNAP (100 µM, 250 µM and 500 µM) or with 1 mM cPTIO. Differences among conditions were analysed using the Kruskal–Wallis test followed by Dunn’s post hoc test with Benjamini– Hochberg correction for multiple comparisons. Different letters indicate statistically significant differences (*P* < 0.05). (N = 141). (B) Time-course transcriptional profiling of PAN. Western blot analysis of PAN transcription levels from 1- to 7-day-old seedlings of *pPAN:GFP* transcriptional reporter line using anti-GFP antibody. Actin protein levels are shown as a loading control. (C) RAM confocal microscopy images of the 5-day-old seedlings of *pPAN:GFP* transcriptional reporter line treated (in the last 6 and 24 hrs at the end of 5 days) with SNAP or cPTIO. Roots were visualised with propidium iodide staining. Bars indicate 20 µm. (D) Boxplots of the corrected total cell fluorescence (CTCF) in QC cells from the 5-day-old seedlings of *pPAN:GFP* transcriptional reporter line treated (in the last 6 and 24 hrs of the end of 5 days) with 0.5 mM SNAP or 1 mM cPTIO. The ANOVA test shows no statistically significant differences (P= 0.662). (N = 51). (E) Western blot analysis of PAN transcription levels in 5-day-old seedlings of *pPAN:GFP* transcriptional reporter line treated with 1 mM SNAP or cPTIO for 5 hrs using anti-GFP antibody. Ponceau S staining showing Rubisco was used as a loading control. (F) Western blot analysis of PAN protein levels in roots from 7-day-old *XVE:PAN* etiolated seedlings using anti-PAN antibody. Seedlings were grown under vertical conditions with 0.5 mM GSNO and 1 mM cPTIO. Actin protein levels are shown as a loading control.

**Supplementary Figure 2 (S2).**
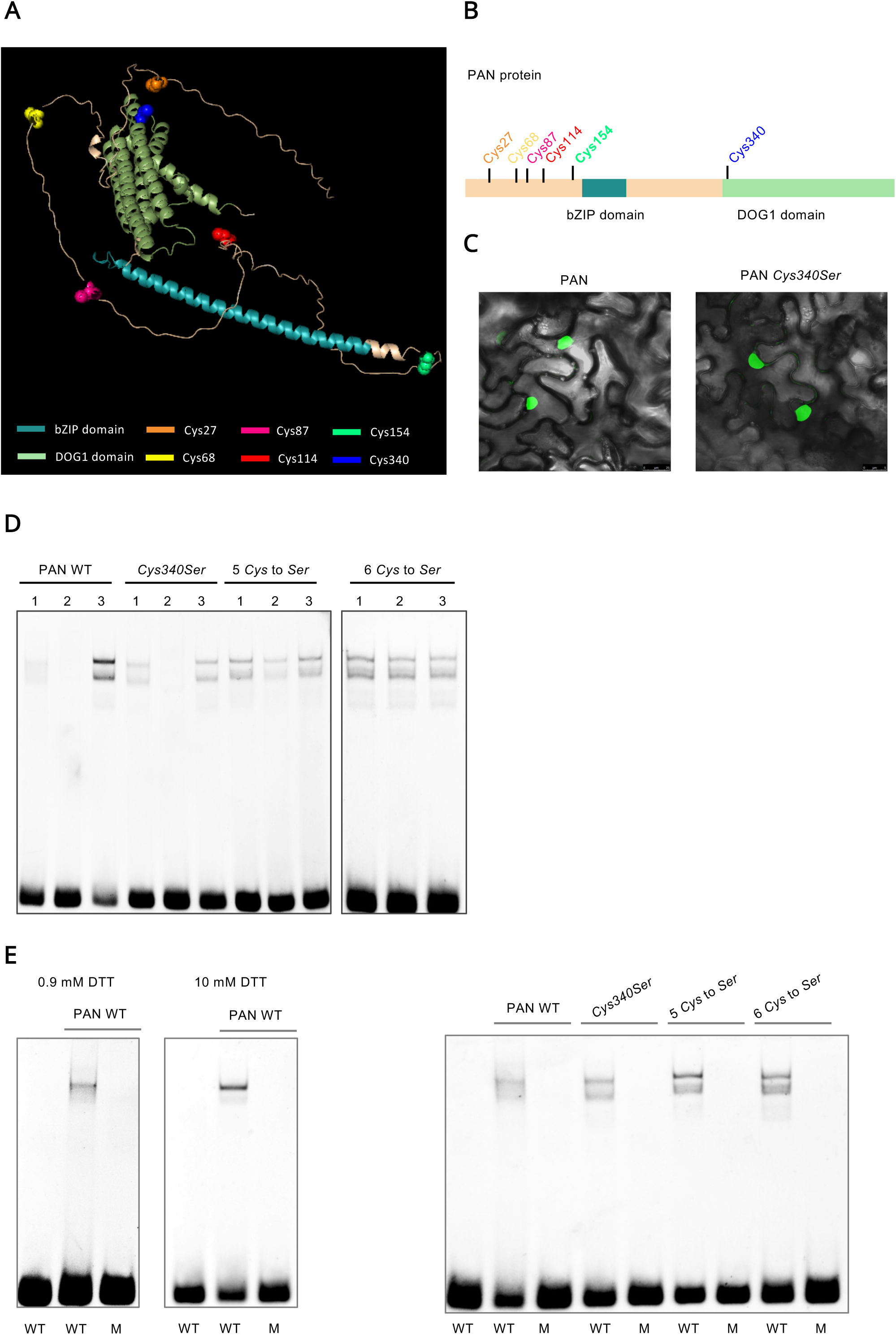
*In silico* structure, subcellular location and DNA binding assays of PAN. *Supporting Figure 2*. (A) 3D structure of PAN predicted with AlphaFold, highlighting the bZIP domain in cyan, the DOG1 domain in green and the Cys residues as sticks. (B) Schematic representation of PAN structure highlighting the bZIP domain in cyan, the DOG1 domain in green and the Cys residues in orange, yellow, pink, red, green and blue. (C) Nuclear localization of WT PAN protein (left) and PAN Cys340Ser mutant protein (right). Agroinfiltration of *Nicotiana benthamiana* leaves with *35S:GFP-PAN* and *35S:GFP-PANC340S* construct, respectively. (D) Electrophoretic Mobility Shift Assay (EMSA) to analyse PAN binding to TGACGT motif. Different PAN versions (WT, *Cys340Ser*, *5 Cys to Ser* and *6 Cys to Ser*) were incubated with 0.9 mM DTT (1), 1 mM GSNO (2) or with 20 mM DTT after treatment with 1 mM GSNO (3). (E) Controls of the Electrophoretic Mobility Shift Assay (EMSA) to analyse PAN binding to TGACGT motif (WT) and mutated version TGATTT (M) under different DTT concentrations in the binding buffer (left) and different PAN versions (WT, *Cys340Ser*, *5 Cys to Ser* and *6 Cys to Ser*) (right).

**Supplementary Figure 3 (S3).**
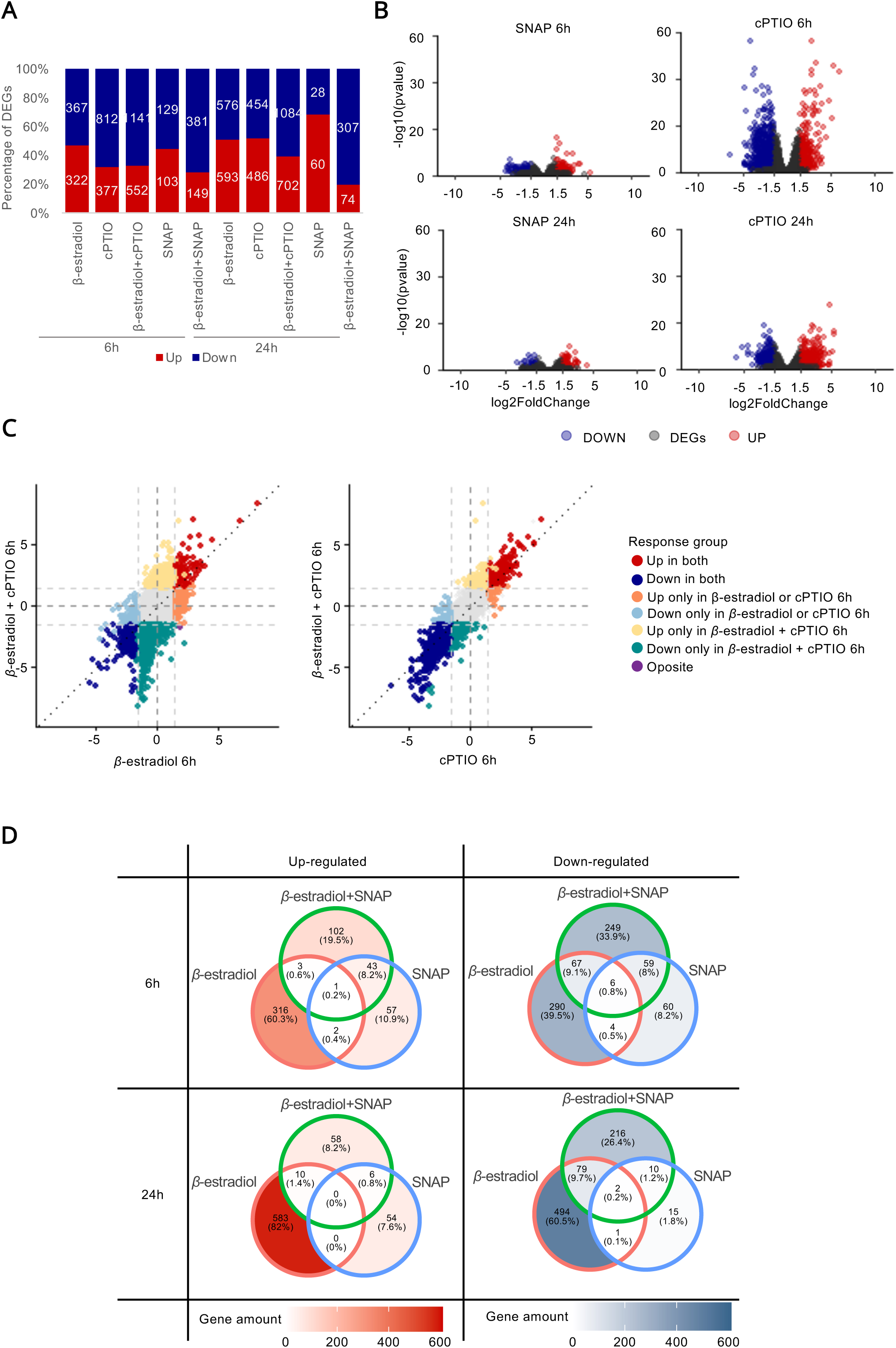
The transcription profile of PAN is co-regulated with NO depletion. *Supporting Figure 3*. (A) Relative frequency of genes up- and down-regulated by condition. Up-regulated DEGs (*qval* ≤ 0.05 and FC ≥ 1.5) are shown in red, while down-regulated DEGs (*qval* ≤ 0.05 and FC ≤ −1.5) are displayed in blue. The numbers within the bars indicate the total gene count. (B) Volcano plots of significant DEGs in the treatments of 0.5 mM SNAP and 1 mM cPTIO during 6 and 24 hours. Red dots depict up-regulated genes (*qval < 0.05* and FC > 1.5), blue ones represent down-regulated genes (*qval < 0.05* and FC < −1.5) and non DEGs (*qval > 0.05)* are coloured grey. (C) Venn diagrams indicating the percentage of common DEGs regulated by SNAP treatments. 6- and 24-hour DEGs are shown separately. Up-regulated DEGs (*qval < 0.05* and FC > 2) are shown in red while down-regulated DEGs (*qval < 0.05* and FC < −2) are displayed in blue. The intensity of the colour correlates with the number of genes.

**Supplementary Figure 4 (S4).**
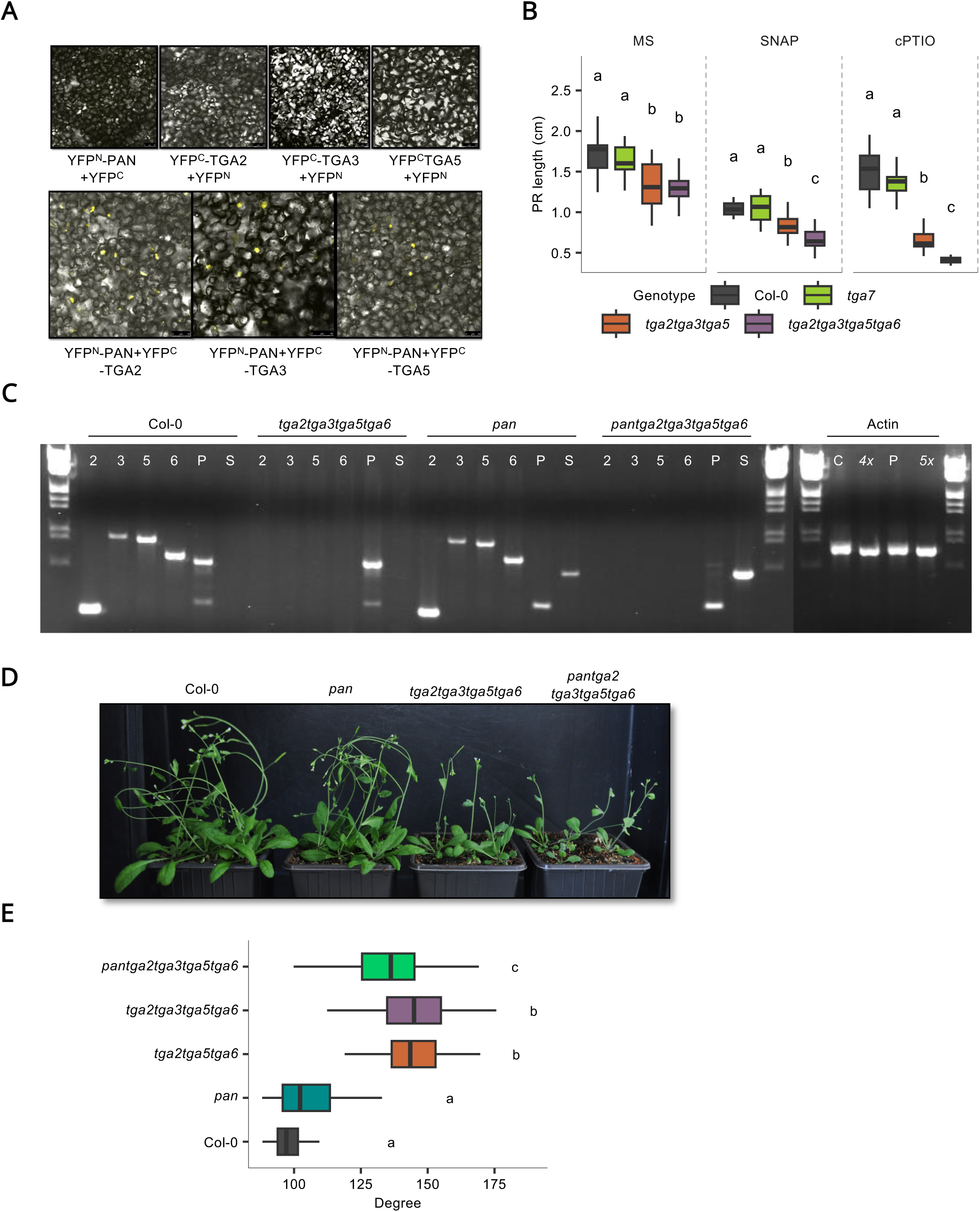
Analysis of the molecular and genetic interaction between PAN and TGA. *Supporting Figure 4*. (A) BiFC assay using agroinfiltration of *Nicotiana benthamiana* leaves to visualise the interaction between PAN and TGA2, TGA3 and TGA5 under a Leica TCS SP2 confocal microscope. Constructs correspond to N-terminal YFP^N^ fusion of PAN (*35S:YFP^N^-PAN*) and N-terminal YFP^C^ fusions of TGA2, TGA3 and TGA5 (*35S:YFP^C^-TGA2,3,5*). (B) Boxplot of PR length of 7-day-old seedlings of Col-0, *tga7*, *triple* and *quadruple* mutants grown on MS, 0.5 mM SNAP or 1 mM cPTIO. Statistical differences among genotypes within each treatment were assessed using a heteroscedastic generalized least-squares model, followed by pairwise comparisons of estimated marginal means with Benjamini–Hochberg adjustment. Different letters indicate statistically significant differences (P<0.05). (N = 232). (C) Genotyping PCR of *pan*, *quadruple* and *pentuple* mutants. *PAN* primers were used for amplification of the 594-bp WT band. *TGA2*, *TGA3*, *TGA5* and *TGA6* primers were used for amplification of the 261-bp, 861-bp, 826-bp and 645-bp bands, respectively. PAN SALK insertion is located 597-bp downstream of the ATG start codon. Actin was used as an endogenous gene control. Primers used for PCR are indicated in Supplementary Table 3. (D) Phenotype of 4-week-old Arabidopsis Col-0, *pan*, *quadruple* and *pentuple* plants. (E) Boxplot of the gravitropic response of *pentuple*, *quadruple* and *pantga2tga5tga6* mutants and Col-0 expressed as the root bending angle after 6 h of gravistimulation. Differences among genotypes were analysed using the Kruskal–Wallis test followed by Dunn’s post hoc test with Benjamini–Hochberg correction for multiple comparisons. Different letters indicate statistically significant differences (*P* < 0.05). (N = 299)

**Supplementary Dataset.**
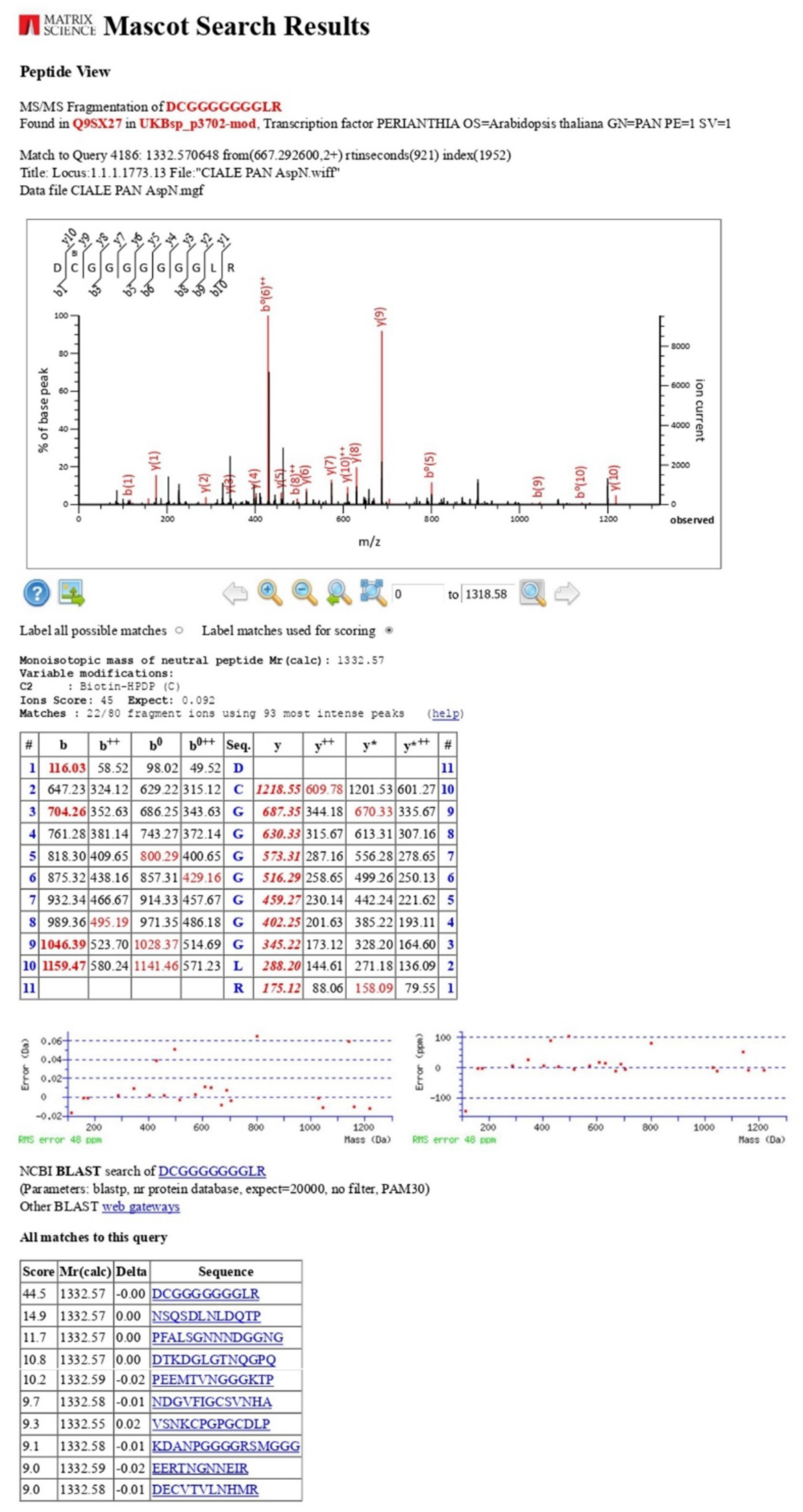
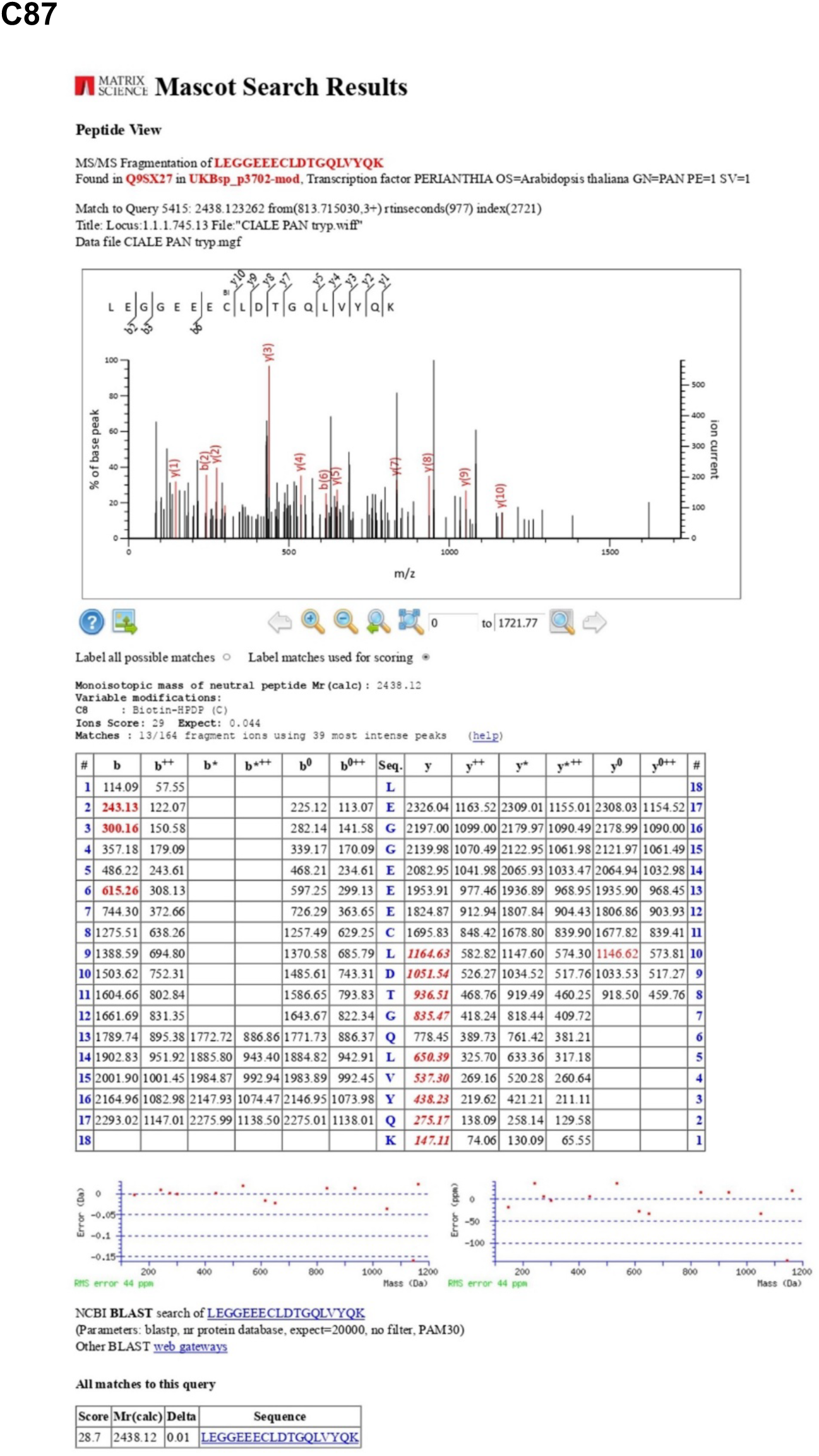
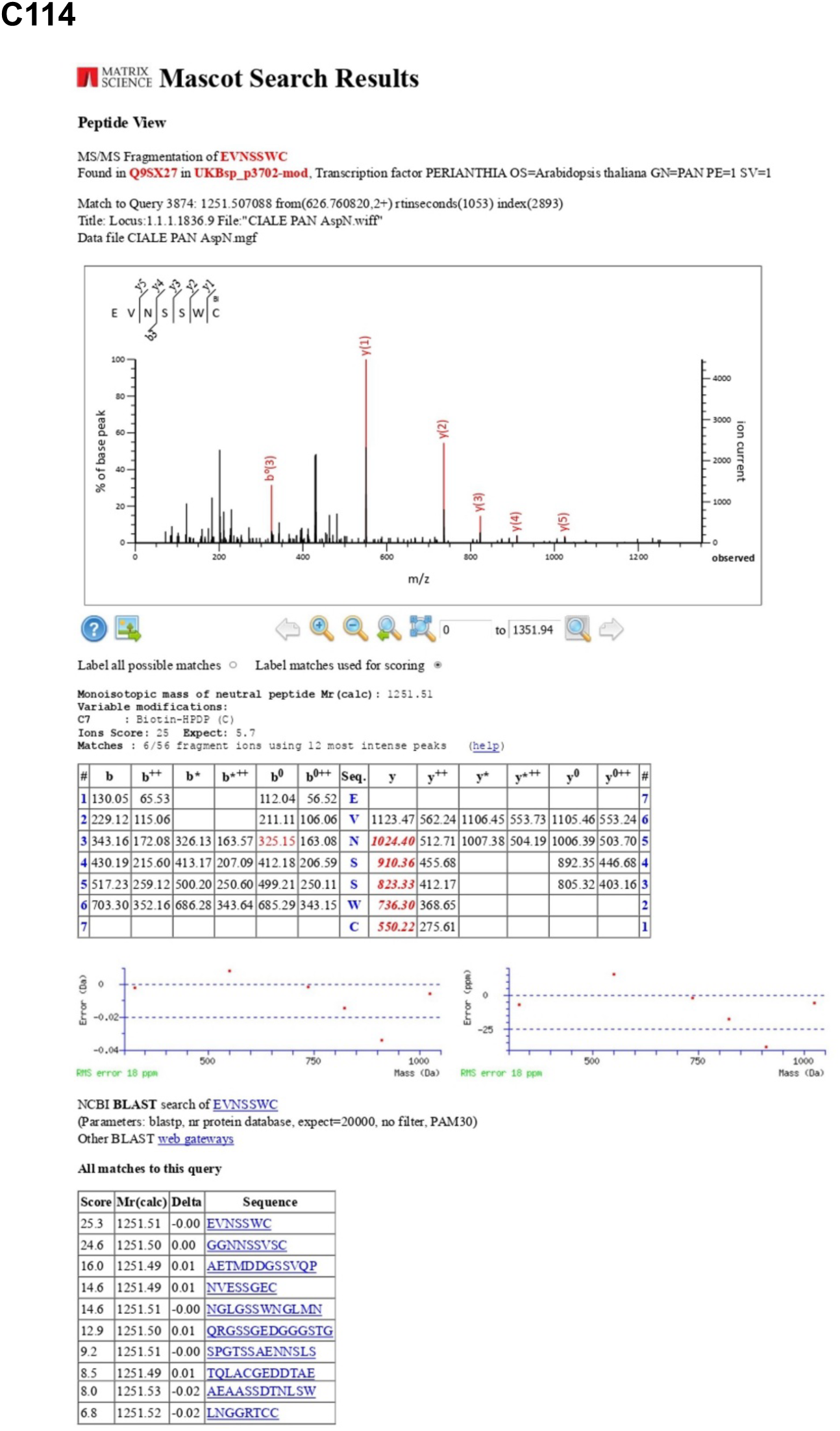
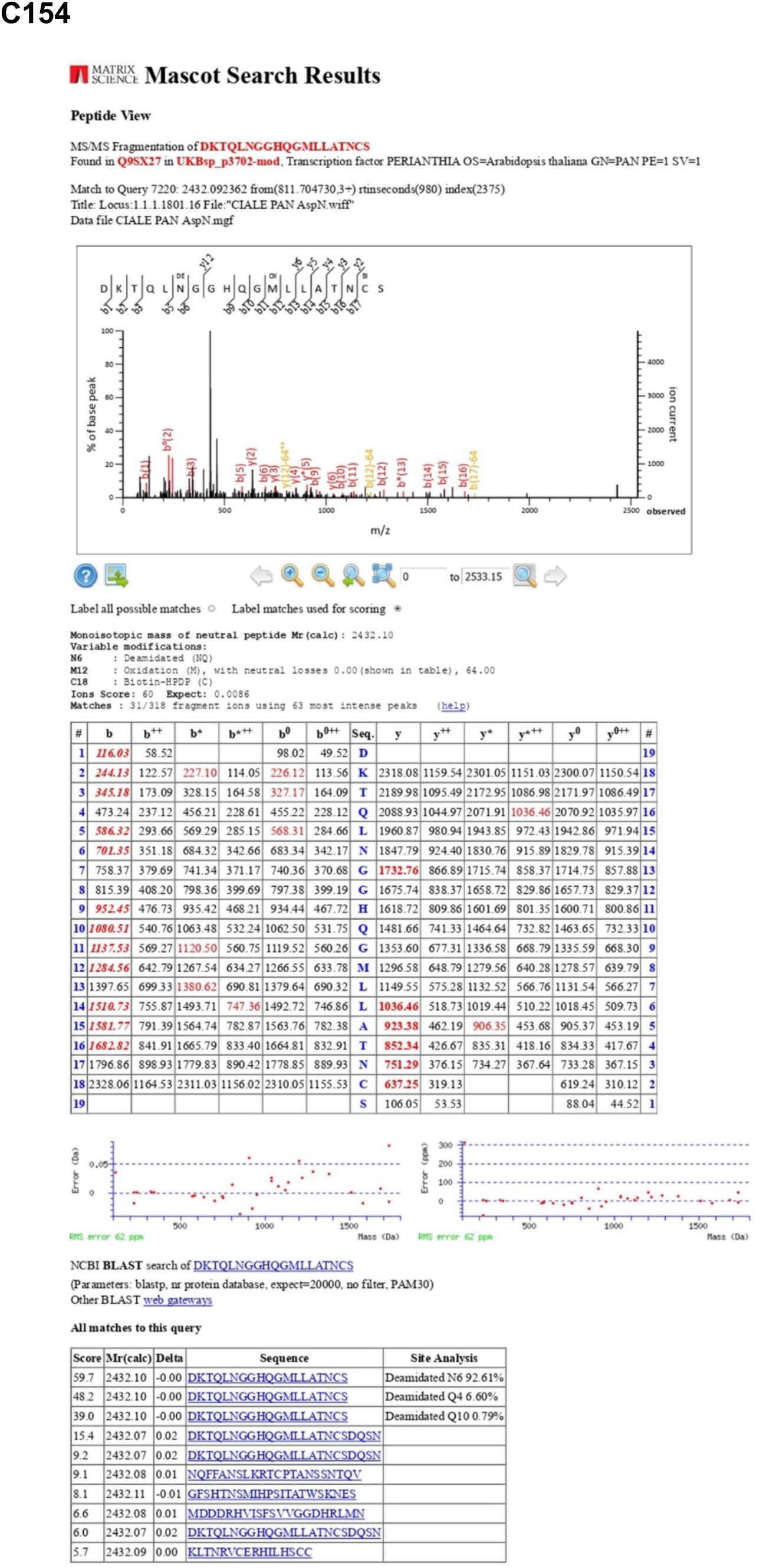
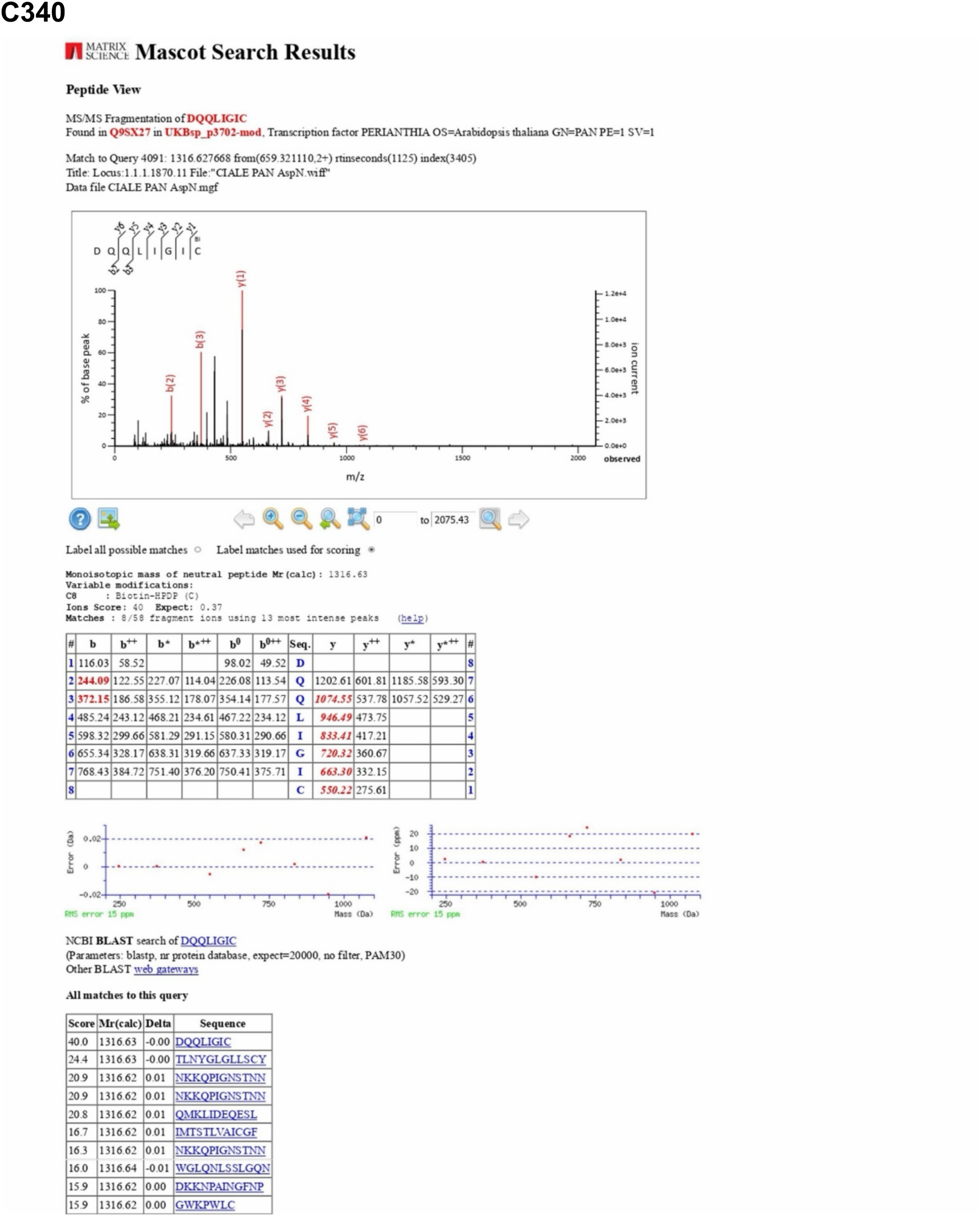
Supporting Figure 2D. Peptide view-MASCOT search showing the MS/MS spectrum and all matched ions, corresponding to the five tryptic fragments in which the Cys residues are modified by Biotin-HPDP, respectively.

## References

Albertos P, Romero-Puertas MC, Tatematsu K, Mateos I, Sánchez-Vicente I, Nambara E, and Lorenzo O. S-nitrosylation triggers ABI5 degradation to promote seed germination and seedling growth. Nat Commun. 2015:6(1):1–10. 10.1038/ncomms9669

Albertos P, Tatematsu K, Mateos I, Sánchez-Vicente I, Fernández-Arbaizar A, Nakabayashi K, Nambara E, Godoy M, Franco JM, Solano R, et al. Redox feedback regulation of ANAC089 signaling alters seed germination and stress response. Cell Rep. 2021:35(11):109263. 10.1016/j.celrep.2021.109263

Alvarez JM, Riveras E, Vidal EA, Gras DE, Contreras-López O, Tamayo KP, Aceituno F, Gómez I, Ruffel S, Lejay L, et al. Systems approach identifies TGA1 and TGA4 transcription factors as important regulatory components of the nitrate response of Arabidopsis thaliana roots. The Plant Journal. 2014:80(1):1–13. 10.1111/TPJ.12618

Andrews SR. FastQC: a quality control tool for high throughput sequence data. 2010.

Astier J and Lindermayr C. Nitric oxide-dependent posttranslational modification in plants: an update. Int J Mol Sci. 2012:13(11):15193–15208. 10.3390/IJMS131115193

Belda-Palazón B, Ruiz L, Martí E, Tárraga S, Tiburcio AF, Culiáñez F, Farràs R, Carrasco P, and Ferrando A. Aminopropyltransferases involved in polyamine biosynthesis localize preferentially in the nucleus of plant cells. PLoS One. 2012:7(10). 10.1371/JOURNAL.PONE.0046907

Belin C, Bashandy T, Cela J, Delorme-Hinoux V, Riondet C, and Reichheld JP. A comprehensive study of thiol reduction gene expression under stress conditions in Arabidopsis thaliana. Plant Cell Environ. 2015:38(2):299–314. 10.1111/PCE.12276/SUPPINFO

Bradford MM. A rapid and sensitive method for the quantitation of microgram quantities of protein utilizing the principle of protein-dye binding. Anal Biochem. 1976:72(1–2):248–254. 10.1016/0003-2697(76)90527-3

Canales J, Contreras-López O, Álvarez JM, and Gutiérrez RA. Nitrate induction of root hair density is mediated by TGA1/TGA4 and CPC transcription factors in Arabidopsis thaliana. Plant Journal. 2017:92(2):305–316. 10.1111/TPJ.13656

Chuang C-F, Running MP, Williams RW, and Meyerowitz EM. The PERIANTHIA gene encodes a bZIP protein involved in the determination of floral organ number in Arabidopsis thaliana. Genes Dev. 1999:13(3):334–344. 10.1101/GAD.13.3.334

Coego A, Brizuela E, Castillejo P, Ruíz S, Koncz C, Del Pozo JC, Piñeiro M, Jarillo JA, León J, Paz-Ares J, et al. The TRANSPLANTA collection of Arabidopsis lines: A resource for functional analysis of transcription factors based on their conditional overexpression. Plant Journal. 2014:77(6):944–953. 10.1111/tpj.12443

Cruz-Ramírez A, Díaz-Triviño S, Blilou I, Grieneisen VA, Sozzani R, Zamioudis C, Miskolczi P, Nieuwland J, Benjamins R, Dhonukshe P, et al. A Bistable Circuit Involving SCARECROW-RETINOBLASTOMA Integrates Cues to Inform Asymmetric Stem Cell Division. Cell. 2012:150(5):1002–1015. 10.1016/J.CELL.2012.07.017

Curtis MD and Grossniklaus U. A Gateway Cloning Vector Set for High-Throughput Functional Analysis of Genes in Planta. Plant Physiol. 2003:133(2):462–469. 10.1104/PP.103.027979

Deblaere R, Bytebier B, de Greve H, Deboeck F, Schell J, van Montagu M, and Leemans J. Efficient octopine Ti plasmid-derived vectors for Agrobacterium-mediated gene transfer to plants. Nucleic Acids Res. 1985:13(13):4777–4788. 10.1093/NAR/13.13.4777

Fernández-Marcos M, Sanz L, Lewis DR, Muday GK, and Lorenzo O. Nitric oxide causes root apical meristem defects and growth inhibition while reducing PIN-FORMED 1 (PIN1)-dependent acropetal auxin transport. Proc Natl Acad Sci U S A. 2011:108(45):18506–18511. 10.1073/pnas.1108644108

Fernández-Marcos M, Sanz L, Lewis DR, Muday GK, and Lorenzo O. Control of Auxin Transport by Reactive Oxygen and Nitrogen Species. In. Polar Auxin Transport., pp. 103–117. 10.1007/978-3-642-35299-7_5

Fernández-Marcos M, Sanz L, and Lorenzo O. Nitric oxide an emerging regulator of cell elongation during primary root growth. Plant Signal Behav. 2012:7(2):196–200. 10.4161/psb.18895

Fox J, Weisberg S, and Price B. Companion to Applied Regression [R package car version 3.1-3]. CRAN: Contributed Packages. 2024. 10.32614/CRAN.PACKAGE.CAR

Gao C-H and Dusa A. ggVennDiagram: A ggplot2 Implement of Venn Diagram. 2024.

Gohel D. rvg: R Graphics Devices for “Office” Vector Graphics Output. 2026.

Graves S, Piepho H-P, and with help from Sundar Dorai-Raj LS. multcompView: Visualizations of Paired Comparisons. 2024.

Gutsche N and Zachgo S. The N-terminus of the floral Arabidopsis TGA transcription factor PERIANTHIA mediates redox-sensitive DNA-binding. PLoS One. 2016:11(4):1–19. 10.1371/journal.pone.0153810

Hess DT, Matsumoto A, Nudelman R, and Stamler JS. S-nitrosylation: spectrum and specificity. Nat Cell Biol. 2001:3(2):E46–E48. 10.1038/35055152

Hothorn T, Bretz F, and Westfall P. Simultaneous Inference in General Parametric Models. Biometrical Journal. 2008:50(3):346–363.

Hu X, Yang L, Ren M, Liu L, Fu J, and Cui H. TGA factors promote plant root growth by modulating redox homeostasis or response. J Integr Plant Biol. 2022:64(8):1543–1559. 10.1111/JIPB.13310

J. DJ. Isolation of plant DNA from fresh tissue. Focus (Madison). 1990:12:13–15.

Kassambara A. ggpubr: ggplot2 Based Publication Ready Plots. 2023a.

Kassambara A. rstatix: Pipe-Friendly Framework for Basic Statistical Tests. 2023b.

Kay M, Elkin LA, Higgins JJ, and Wobbrock JO. ARTool: Aligned Rank Transform. 2021.

Kim D, Paggi JM, Park C, Bennett C, and Salzberg SL. Graph-based genome alignment and genotyping with HISAT2 and HISAT-genotype. Nat Biotechnol. 2019:37(8):907–915. 10.1038/s41587-019-0201-4

Kneeshaw S, Gelineau S, Tada Y, Loake GJ, and Spoel SH. Selective protein denitrosylation activity of Thioredoxin-h5 modulates plant Immunity. Mol Cell. 2014:56(1):153–162. 10.1016/J.MOLCEL.2014.08.003

Kolde R. Pheatmap: Pretty Heatmaps. R package version 1.0.12. 2019.

Kovacs I and Lindermayr C. Nitric oxide-based protein modification: formation and site-specificity of protein S-nitrosylation. Front Plant Sci. 2013:4(137):1–10. 10.3389/FPLS.2013.00137

Lee J-Y, Colinas J, Wang JY, Mace D, Ohler U, and Benfey PN. Transcriptional and posttranscriptional regulation of transcription factor expression in Arabidopsis roots. PNAS. 2006:103(15):6055–6060.

Van Leene J, Boruc J, De Jaeger G, Russinova E, and De Veylder L. A kaleidoscopic view of the Arabidopsis core cell cycle interactome. Trends Plant Sci. 2011:16(3):141–150. 10.1016/J.TPLANTS.2010.12.004

Van Leene J, Eeckhout D, Cannoot B, De Winne N, Persiau G, Van De Slijke E, Vercruysse L, Dedecker M, Verkest A, Vandepoele K, et al. An improved toolbox to unravel the plant cellular machinery by tandem affinity purification of Arabidopsis protein complexes. Nat Protoc. 2015:10(1):169–187. 10.1038/nprot.2014.199

Lenth R V and Piaskowski J. emmeans: Estimated Marginal Means, aka Least-Squares Means. 2026.

Li M, Yao T, Lin W, Hinckley WE, Galli M, Muchero W, Gallavotti A, Chen JG, and Huang S shan C. Double DAP-seq uncovered synergistic DNA binding of interacting bZIP transcription factors. Nature Communications 2023 14:1. 2023:14(1):1–19. 10.1038/s41467-023-38096-2

Li S, Lauri A, Ziemann M, Busch A, Bhave M, and Zachgo S. Nuclear activity of ROXY1, a glutaredoxin interacting with TGA factors, is required for petal development in Arabidopsis thaliana. Plant Cell. 2009:21(2):429–441. 10.1105/tpc.108.064477

Li S, Yamada M, Han X, Ohler U, and Benfey PN. High-Resolution Expression Map of the Arabidopsis Root Reveals Alternative Splicing and lincRNA Regulation. Dev Cell. 2016:39(4):508–522. 10.1016/J.DEVCEL.2016.10.012

Lindermayr C, Sell S, Müller B, Leister D, and Durner J. Redox Regulation of the NPR1-TGA1 System of Arabidopsis thaliana by Nitric Oxide. Plant Cell. 2010:22(8):2894–2907. 10.1105/TPC.109.066464

Lorenzo O. BZIP edgetic mutations: At the frontier of plant metabolism, development and stress trade-off. J Exp Bot. 2019:70(20):5517–5520. 10.1093/jxb/erz298

Love MI, Anders S, and Huber W. Differential analysis of count data - the DESeq2 package. Genome Biol. 2014:15(12):550. 10.1186/s13059-014-0550-8

de Luis Balaguer MA, Fisher AP, Clark NM, Fernandez-Espinosa MG, Möller BK, Weijers D, Lohmann JU, Williams C, Lorenzo O, and Sozzani R. Predicting gene regulatory networks by combining spatial and temporal gene expression data in Arabidopsis root stem cells. Proc Natl Acad Sci U S A. 2017:114(36):E7632–E7640. 10.1073/pnas.1707566114

Maier AT, Stehling-Sun S, Offenburger SL, and Lohmann JU. The bZIP Transcription Factor PERIANTHIA: A Multifunctional Hub for Meristem Control. Front Plant Sci. 2011:2(79):1–17. 10.3389/FPLS.2011.00079

Maier AT, Stehling-Sun S, Wollmann H, Demar M, Hong RL, Haubeiß S, Weigel D, and Lohmann JU. Dual roles of the bZIP transcription factor PERIANTHIA in the control of floral architecture and homeotic gene expression. Development. 2009:136(10):1613–1620. 10.1242/dev.033647

Mata-Pérez C, Sánchez-Vicente I, Arteaga N, Gómez-Jiménez S, Fuentes-Terrón A, Oulebsir CS, Calvo-Polanco M, Oliver C, and Lorenzo Ó. Functions of nitric oxide-mediated post-translational modifications under abiotic stress. Front Plant Sci. 2023:14(1158184):1–15. 10.3389/FPLS.2023.1158184/BIBTEX

Millard SP. EnvStats: an R package for environmental statistics. 2013:291.

Nakagawa T, Ishiguro S, and Kimura T. Gateway vectors for plant transformation Invited Review.

Nawy T, Lee JY, Colinas J, Wang JY, Thongrod SC, Malamy JE, Birnbaum K, and Benfey PN. Transcriptional Profile of the Arabidopsis Root Quiescent Center. Plant Cell. 2005:17(7):1908–1925. 10.1105/TPC.105.031724

Pavelescu I, Vilarrasa-Blasi J, Planas-Riverola A, González-García M, Caño-Delgado AI, and Ibañes M. A Sizer model for cell differentiation in Arabidopsis thaliana root growth. Mol Syst Biol. 2018:14(1):e7687. 10.15252/MSB.20177687/SUPPL_FILE/MSB177687-SUP-0010-DATASETEV2.ZIP

Pinheiro J, Bates D, and R Core Team. nlme: Linear and Nonlinear Mixed Effects Models. 2024.

R Core Team. R: A Language and Environment for Statistical Computing. 2025.

Sambrook J, Fritsch EF, and Maniatis T. Molecular cloning: a laboratory manual. 2nd ed.

Sanchez-Corrionero A, Sánchez-Vicente I, Arteaga N, Manrique-Gil I, Gómez-Jiménez S, Torres-Quezada I, Albertos P, and Lorenzo O. Fine-Tuned Nitric Oxide And Hormone Interface In Plant Root Development And Regeneration. J Exp Bot. 2023:74(19):6104–6118. 10.1093/JXB/ERAC508

Sánchez-Vicente I, Albertos P, Sanz C, Wybouw B, De Rybel B, Begara-Morales JC, Chaki M, Mata-Pérez C, Barroso JB, and Lorenzo O. Reversible S-nitrosylation of bZIP67 by peroxiredoxin IIE activity and nitro-fatty acids regulates the plant lipid profile. Cell Rep. 2024:43(4):114091. 10.1016/J.CELREP.2024.114091

Sánchez-Vicente I, Fernández-Espinosa MG, and Lorenzo O. Nitric oxide molecular targets: Reprogramming plant development upon stress. J Exp Bot. 2019:70(17):4441–4460. 10.1093/jxb/erz339

Sanz L, Albertos P, Mateos I, Sánchez-Vicente I, Lechón T, Fernández-Marcos M, and Lorenzo O. Nitric oxide (NO) and phytohormones crosstalk during early plant development. J Exp Bot. 2015:66(10):2857–2868. 10.1093/jxb/erv213

Sanz L, Fernández-Marcos M, Modrego A, Lewis DR, Muday GK, Pollmann S, Dueñas M, Santos-Buelga C, and Lorenzo O. Nitric oxide plays a role in stem cell niche homeostasis through its interaction with auxin. Plant Physiol. 2014:166(4):1972–1984. 10.1104/pp.114.247445

Sato EM, Hijazi H, Bennett MJ, Vissenberg K, and Swarup R. New insights into root gravitropic signalling. J Exp Bot. 2015:66(8):2155–2165. 10.1093/JXB/ERU515

Schindelin J, Arganda-Carreras I, Frise E, Kaynig V, Longair M, Pietzsch T, Preibisch S, Rueden C, Saalfeld S, Schmid B, et al. Fiji: an open-source platform for biological-image analysis. Nat Methods. 2012:9(7):676–682. 10.1038/nmeth.2019

Truernit E, Bauby H, Dubreucq B, Grandjean O, Runions J, Barthélémy J, and Palauqui JC. High-Resolution Whole-Mount Imaging of Three-Dimensional Tissue Organization and Gene Expression Enables the Study of Phloem Development and Structure in Arabidopsis. Plant Cell. 2008:20(6):1494–1503. 10.1105/TPC.107.056069

Wickham H. ggplot2: Elegant Graphics for Data Analysis. 2016.

Van Der Zaal EJ, Droog FNJ, Boot CJM, Hensgens LAM, Hoge JHC, Schilperoort RA, and Libbenga KR. Promoters of auxin-induced genes from tobacco can lead to auxin-inducible and root tip-specific expression. Plant Mol Biol. 1991:16(6):983–998.

